# A hierarchical immune receptor network in lettuce reveals contrasting patterns of evolution in sensor and helper NLRs

**DOI:** 10.1101/2025.02.25.639832

**Authors:** Hsuan Pai, Toshiyuki Sakai, Andres Posbeyikian, Raoul Frijters, Yu Sugihara, Mauricio P. Contreras, Jiorgos Kourelis, Hiroaki Adachi, Sophien Kamoun, AmirAli Toghani

## Abstract

Nucleotide-binding domain and leucine-rich repeat immune receptors (NLRs) are known for their rapid evolution, even at the intraspecific level, yet the rates of evolution differ significantly across various NLR classes. Within the NRC (NLR Required for Cell Death) network, NLRs operate in complex sensor-helper configurations to confer immunity against a diverse array of pathogens, particularly in Asterids. While helper NLRs are typically conserved and evolve slowly, sensor NLRs tend to evolve more rapidly. However, the functional connections between slow and fast-evolving NLRs remain poorly understood, notably in important crop species. We conducted a comparative analysis of NLRs across 40 Solanales and 29 Asterales genomes to explore NRC network expansion and diversification within the less-studied Asterales order. Our findings reveal that the NRC network has expanded less in Asterales compared to Solanales. We functionally validated a minimal Asterales NRC network with 2 helpers and 9 sensors in common lettuce (*Lactuca sativa*). Through selection and diversification analysis and structural modeling of NRC helper and sensor subclades in the *Lactuca* genus, we found varying evolutionary diversification rates between NRC helpers and sensors. We found a correlation between sensor diversification rates and helper dependency, with sensors reliant on a phylogenetically conserved helpers experiencing limited diversification pressure. Our results highlight the lineage- and function-specific evolution of the NRC network, offering insights into the evolutionary pressures shaping plant immune receptor networks.

## INTRODUCTION

Plants possess a sophisticated innate immune system to defend against a wide array of pathogens (Jones and Dangl 2006; Kourelis and van der Hoorn 2018). A group of intracellular immune receptors, known as nucleotide-binding domain and leucine-rich repeat-containing receptors (NLRs), are essential components of this defense system. These receptors detect pathogen-secreted effector proteins. Upon recognition, NLRs trigger a cascade of defense responses, often culminating in a localized programmed cell death that effectively restricts pathogen spread (Jones et al. 2016; Contreras et al. 2023a). NLR proteins represent one of the most diverse protein families, with different NLR clades evolving at varying rates (Van de Weyer et al. 2019; Prigozhin and Krasileva 2021). However, the relationship between the rate of NLR evolution and their functional roles remains poorly understood.

NLR proteins are typically composed of three main domains: an N-terminal domain, often a coiled-coil (CC) or Toll-interleukin receptor (TIR), which is responsible for signal transduction; a central NB-ARC (nucleotide-binding domain shared with APAF-1, plant R proteins, and CED-4) domain which and plays a role in regulating activation and signaling; and a C-terminal leucine-rich repeat (LRR) domain, which directly or indirectly facilitates pathogen detection (Kourelis and van der Hoorn 2018; Duxbury et al. 2021; Kourelis et al. 2021). Based on their NB-ARC phylogeny, NLRs are classified into four subclades: CC-NLRs (CNLs), CC_G10_-type CC-NLRs, RPW8-type CC-NLRs (CC_R_-NLRs/RNLs), and TIR-NLRs (TNLs) (Kourelis et al. 2021; Lee et al. 2021). While the tripartite architecture is largely conserved, some NLRs contain additional or non-canonical domains, such as integrated domains (IDs), which contribute to pathogen detection (Cesari et al. 2014; Maqbool et al. 2015; Baggs et al. 2017; Grund et al. 2019). About 20% of CC-NLRs harbor a conserved N-terminal MADA motif essential for triggering cell death (Adachi et al. 2019a). Activated CC-NLRs often form higher-order complexes called “resistosome” (Wang et al. 2019; Förderer et al. 2022; Zhao et al. 2022; Liu et al. 2024; Madhuprakash et al. 2024), which likely insert into the plasma membrane to act as calcium channels via their N-terminal alpha-helix (Wang et al. 2019; Bi et al. 2021; Jacob et al. 2021). Recent advances in computational structural biology have allowed for modeling NLR resistosomes in the presence of simulated plasma membrane with high confidence (Madhuprakash et al. 2024; Toghani et al. 2024).

While some NLRs function as independent units, many form pairs or networks of specialized NLRs to enhance evolvability and robustness of immune responses (Wu et al. 2018; Contreras et al. 2023a). NLR pairs, often clustered genetically, work together, such as Pik-1/Pik-2 and RGA5/RGA4 CC-NLR pairs in rice (*Oryza sativa*) and the RPS4/RRS1 TIR-NLR pair in *Arabidopsis thaliana* (Narusaka et al. 2009; Cesari et al. 2014; Le Roux et al. 2015; Maqbool et al. 2015; Sarris et al. 2015; Shimizu et al. 2022; Sugihara et al. 2023). In NLR networks, sensor NLRs depend on helper NLRs for immunity, following one-to-many or many-to-one configurations (Wu et al. 2017, 2018; Adachi et al. 2019b; Contreras et al. 2023a). For instance, the CC_R_-NLRs NRG1 and ADR1 can function downstream of the TIR-NLR signaling pathway (Qi et al. 2018; Castel et al. 2019; Lapin et al. 2019; Saile et al. 2020). The NRC network, first identified in Solanaceae plants like tomato and *Nicotiana benthamiana*, also exemplifies a CC-NLR network, where sensor NLRs rely on NRC helpers (NRC-H) to activate immune responses (Wu et al. 2017, 2018). NRC-dependent sensors (NRC-S) are divided into two subtypes based on domain architecture: Rx-type sensors have a canonical CC-NB-ARC-LRR structure, while SD-type sensors include an additional N-terminal Solanaceous Domain (SD) that contributes to pathogen recognition (Li et al. 2019; Seong et al. 2020; Contreras et al. 2023a; Lüdke et al. 2023). Upon pathogen effector detection, NRC sensors transmit a signal that triggers the assembly of helper NRCs from a homodimer resting state into hexameric resistosomes that induce immune responses and cell death (Contreras et al. 2022; Ahn et al. 2023; Liu et al. 2024; Madhuprakash et al. 2024; Selvaraj et al. 2024). Though the exact mechanism of signal transmission between sensor and helper NRCs remains unclear, it likely involves transient “activation-and-release” interactions (Contreras et al. 2022; Ahn et al. 2023).

The NRC network superclade, consisting of NRC-H and NRC-S, likely originated ∼100 million years ago within the superasterid lineage of land plants and has expanded massively in the lamiid lineage (Wu et al. 2017; Goh et al. 2024; Sakai et al. 2024). While particularly prominent in Solanaceae, its expansion and complexity vary across plant families (Goh et al. 2024). *NRC0*, the most conserved and ancient NRC helper in asterids, represents the ancestral state of the NRC network and is frequently genetically linked to its dependent NRC-S (Goh et al. 2024; Sakai et al. 2024). In contrast, other NRC subclades have undergone lineage-specific expansions and exhibit diverse genomic arrangements, with some NRC-H clustered alongside NRC-S, such as *NRC6*, and others dispersed throughout the genome (Wu et al. 2020; Lüdke et al. 2023; Goh et al. 2024). While sensor NLRs evolve rapidly to counter pathogen evolution, helper NLRs are typically more conserved (Goh et al. 2024). However, recent findings show orthologous helpers also experience diversification pressures to prevent cross-activation and isolate signaling pathways (Selvaraj et al. 2024).

NLR numbers vary widely across species, from a few to over a thousand in some species (Baggs et al. 2020; Lee and Chae 2020; Liu et al. 2021; Goh et al. 2024; Toghani and Kamoun 2024), reflecting the dynamic co-evolutionary arms race between plants and pathogens (Barragan and Weigel 2021). NLR gene evolution follows the birth-and-death model, where gene duplication generates new NLRs—some are maintained and adapt to detect pathogens, while others lose function or are deleted (Michelmore and Meyers 1998; Seong et al. 2020; Barragan and Weigel 2021; Cruz et al. 2024). Recent advances have introduced new evolutionary models, emphasizing the dynamic response of NLRs to pathogen pressure (Barragan and Weigel 2021; Shimizu et al. 2022; Cruz et al. 2024; Sugihara et al. 2024). Pathogen effectors often target NLRs to suppress immunity, but NLRs counteract this through mutations in critical regions or by developing functional redundancy (Wu et al. 2018; Sugihara et al. 2024). For instance, within the NRC network, helpers like NRC2 and NRC3 are suppressed by cyst nematode effector SS15 and oomycete effector AVRcap1b (Derevnina et al. 2021; Contreras et al. 2023b). However, in some species NRC3 has accumulated mutations in its effector-binding region to evade suppression (Sugihara et al. 2024). Additionally, signaling redundancy allows non-suppressed nodes, such as NRC4, to activate immune responses even when parts of the pathway are suppressed (Wu et al. 2018; Contreras et al. 2023b; Huang et al. 2023; Sugihara et al. 2024).

Asterales is one of the most diverse flowering plant orders within the Asterid clade, encompassing several commercially important crops, including the common lettuce (*Lactuca sativa*) (Collinson et al. 2000; Smith and Brown 2018). Lettuce has become a key model for studying genetic resistance to pathogens and NLR evolution (Meyers et al. 1998; Shen et al. 2002; McHale et al. 2009; Truco et al. 2013; Christopoulou et al. 2015; Parra et al. 2016, 2021b, 2021a; Sandoya et al. 2021; Pink et al. 2022). Meyers et al. (1998) categorized the lettuce NLRome into 42 phylogenetic clades, known as Resistance Gene Clusters (RGCs) (Meyers et al. 1998; McHale et al. 2009). These NLR genes were further grouped into eight Major Resistance Clusters (MRCs) based on their locations on chromosomes 1, 2, 3, 4, 8, and 9 (McHale et al. 2009; Truco et al. 2013). In another study, Kuang et al. (2004) reported that genes at the RGC2 locus in lettuce could be categorized into two types: type I (diverse) and type II (conserved), with contrasting patterns of evolution and copy number between them (Kuang et al. 2004). This concept has since been explored in other systems and NLR families, including *Arabidopsis thaliana* and *Zea mays* (maize) (Prigozhin and Krasileva 2021; Prigozhin et al. 2024; Sutherland et al. 2024). Despite extensive functional and genetic research on resistance in lettuce, the evolution of the NRC network in Asterales remain poorly understood.

In this study, we analyzed 40 Solanales and 29 Asterales genomes to investigate the expansion and diversification of the NRC network within the Asterales order. We found that the NRC network in Asterales shows limited expansion compared to Solanales. Notably, NLRs in the Rx-type sensor phylogenetic clade are absent in Asterales, and although NRC-S in Asterales are phylogenetically classified as SD-types based on their NB-ARC, they lack the N-terminal Solanaceous domain. We also confirmed the presence of a functional NRC network in common lettuce (*Lactuca sativa*). Through selection and diversification analyses of NRC helper and sensor subclades within the *Lactuca* genus, we observed distinct patterns of diversification—not only between helpers and sensors but also among different sensor clades. Our results suggest that NRC sensors dependent on the conserved helper NRC0 show limited diversification and expansion, whereas sensors relying on other NRC helpers have undergone greater expansion and display higher rates of diversification. We further showed that unlike NRC helpers, NRC sensors are not predicted to form resistosome-like structures when modeled with AlphaFold 3. These findings provide insights into NRC network evolution and the functional relationships among differentially evolving NLR clades.

## RESULTS

### The NRC network superclade shows limited expansion in Asterales compared to Solanales

The presence of NLRs in the NRC network superclade across Asterids prompted us to investigate the extent of its expansion in Asterales, a Campanulid order, compared to the well-studied Solanales order from Lamiids (**Figure 1A**) (Wu et al. 2017). For this purpose, we extracted a total number of 21,232 non-redundant NLR sequences from a dataset of de-novo annotated 40 Solanales and 29 Asterales genomes with harmonized annotations, representing a diverse set of species and genera from both orders (**Figure S1**, **Data S1 and S2**) (Toghani et al. 2025a). Based on phylogenetics analysis we then extracted the monophyletic NRC network superclade comprising 7,020 sequences, including the reference NRC-H and NRC-S from RefPlantNLR (**Data S3**) (Kourelis et al. 2021). In Asterales and Solanales, there are on average 300 and 338.6 non-redundant NLRs per genome, respectively. However, NRC superclade NLRs on average only comprise 6.6% of the NLRome in Asterales genomes compared to 50.3% in Solanales (**Figure S2**).

**Figure 1.**
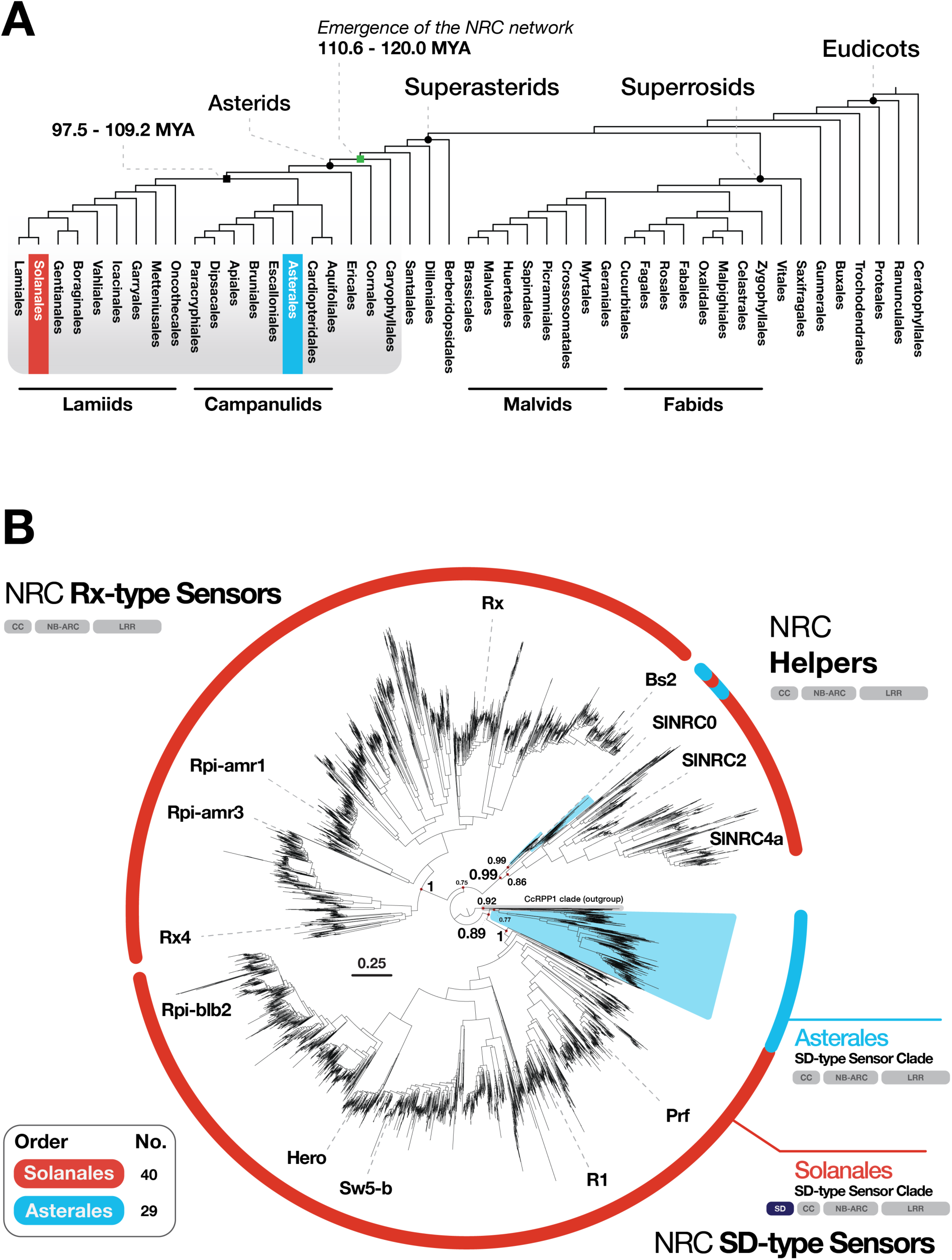
The NRC network superclade has expanded less in Asterales compared to Solanales. (A) Taxonomic tree of eudicots with NRC network superclade timeline in Asterids. Phylogenetic tree of plant orders and divergence time estimates obtained from TimeTree 5 (Kumar et al. 2022). (B) Phylogenetics tree of 7,040 NRC network superclade and CcRPP1 paraphyletic clade sequences from 40 Solanales and 29 Asterales genomes. Asterales’ clades and branches are highlighted in blue.

Compared to Solanales, Asterales show a significantly less expanded NRC network, as indicated by the lower copy numbers in both sensor and helper clades (**Figure 1B and S2**). We divided the 88 Asterales NRC helpers into two subclades. One subclade, containing 27 sequences clustered with the conserved NRC0 clade, while the remaining 61 sequences formed a separate clade and clustered with known Solanales NRC helpers, such as NRC2 and NRC4, referred to as Ast-NRCs hereafter (**Figure 1B and S2**) (Wu et al. 2017). All 504 Asterales NRC sensors formed a distinct, well-supported clade that grouped with the Solanales SD-type sensors, but none are phylogenetically clustered with the Rx-type Solanales sensor clade (**Figure 1B**). Although Asterales NRC sensors belong to the SD-type clade, they lack the Solanaceous domain characteristic at their N-termini (**Figure 1B, Data S3**) (Seong et al. 2020). These observations suggest a substantial difference in the evolutionary pace and expansion of the NRC network between Solanales and Asterales orders.

In common lettuce, the 17 NRC superclade sequences (comprising 2 helpers and 15 sensors) represent only 5% of the total NLRome, which consists of 359 sequences. In contrast, TIR-NLRs dominate the NLRome with 206 sequences, making up over 57%. The CC-NLR, CC_R_-NLR, and CC_G10_-NLR clades contain 97, 9, and 42 sequences, respectively (**Figure S3**). This indicates a disproportionate expansion of different NLR types, with a limited expansion of NRC-H and NRC-S in lettuce.

### NRC helpers and sensors are often genetically clustered in lettuce and other Asterales species

Given the observed differences in the expansion of NRC network superclade between Solanales and Asterales, we further investigated the Asterales NRC network superclade in more detail, focusing on lettuce. We analyzed 40 NRC-H and NRC-S sequences from common lettuce (*Lactuca sativa*) and two wild lettuce species (*Lactuca saligna* and *Lactuca virosa*) (**Data S4**). Phylogenetic analysis revealed four subclades in the *Lactuca* NRC superclade sequences: one helper clade containing sequences that grouped with either NRC0 or tomato NRC2 (referred to as Ast-NRC1 in each species hereafter), and sensor clades 1, 2, and 3 (**Figure 2A**). These subclades were defined by well-supported branches that form distinct clades in the phylogenetic tree (**Figure 2A and S5**). Based on the previous categorization of NLR families in lettuce, NRC superclade clade sequences are distributed across four RGC groups: (1) the NRC helper clade as RGC7, (2) sensor clade 1 as RGC26, (3) sensor clade 2 as RGC27, and (4) sensor clade 3 as RGC9 (**Figure S3**) (Meyers et al. 1998; McHale et al. 2009).

**Figure 2.**
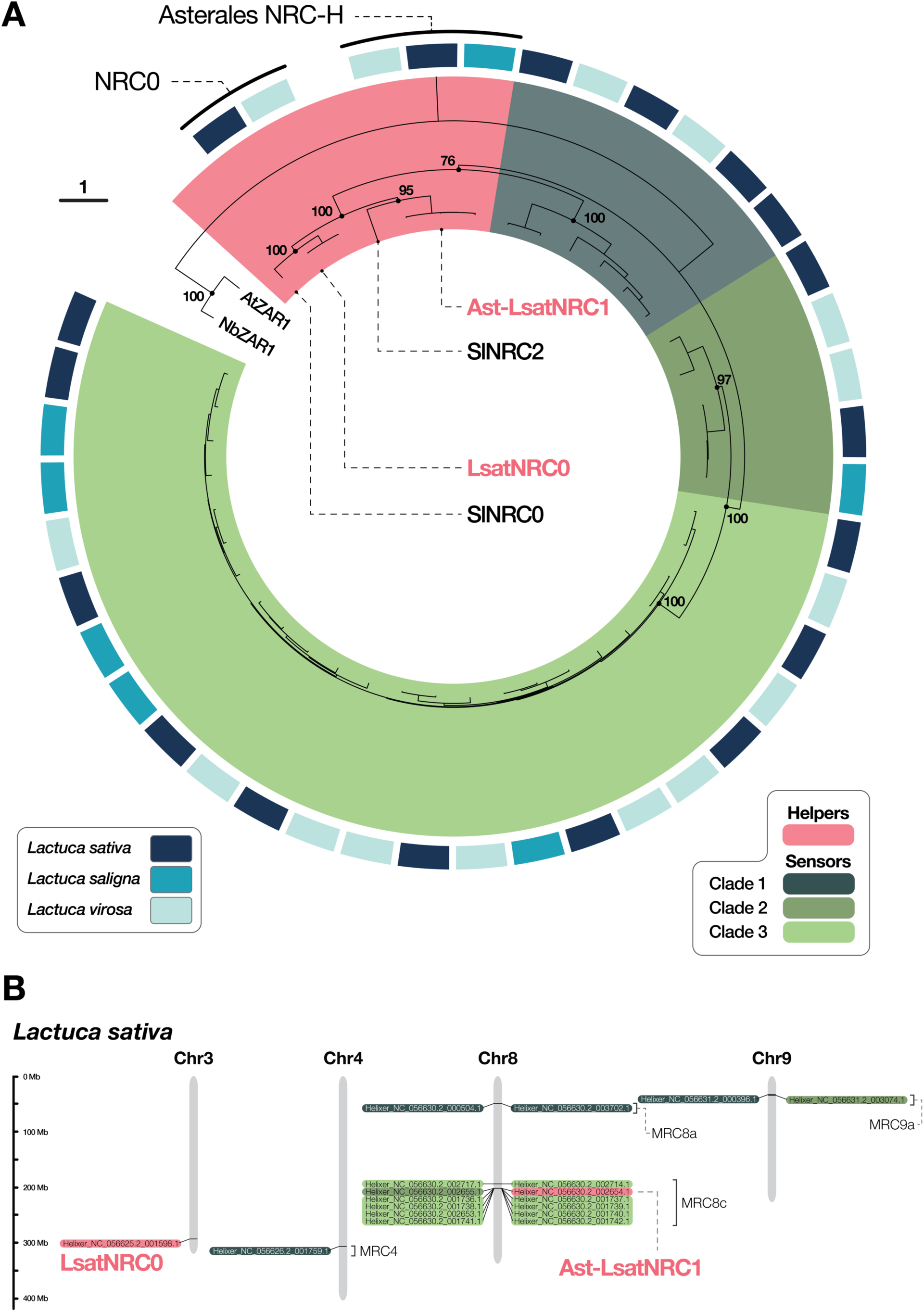
Unlike LsatNRC0, Ast-LsatNRC1 is genetically linked to sensor NLRs in lettuce. (A) Phylogenetic tree of NRC superclade sequences of three *Lactuca* species (*Lactuca sativa*, *Lactuca saligna*, and *Lactuca virosa*). NRC-S are divided in three well-supported phylogenetic subclades. Numbers on tree nodes indicate bootstrap values. AtZAR1 and NbZAR1 were used as outgroup sequences. SlNRC0 and SlNRC2 were used as reference sequences. (B) Physical map of *Lactuca sativa* NRC superclade sequences on the chromosomes. Previously described Major Resistance Clusters (MRCs) are shown for NRCs (McHale et al. 2009; Truco et al. 2013).

Based on previous reports of NRC-H and NRC-S being in genetic clusters for NRC0 and NRC6 (Lüdke et al. 2023; Sakai et al. 2024), we examined the physical location of NRC superclade sequences in the lettuce genomes. Unlike previous observations in various Asterid species, lettuce NRC0 (LsatNRC0; Helixer_NC_056625.2_001598.1), on chromosome 3, was not located near any sensor sequences (**Figure 2B**) (Sakai et al. 2024). In contrast, Ast-LsatNRC1 (Helixer_NC_056630.2_002654.1) was found to be in a physical cluster (<100kb) with sensors from clades 2 and 3, all residing on chromosome 8. Notably, all of clade 3 sensors are clustered together on this chromosome. This cluster is part of the previously reported MRC8c (**Figure 2B**) (Meyers et al. 1998; McHale et al. 2009; Christopoulou et al. 2015). The remaining lettuce NRC-S sequences, including those from clade 1, are scattered across chromosomes 4, 8, and 9, as part of MRCs 4, 8a, and 9a, respectively (**Figure 2B**) (Meyers et al. 1998; McHale et al. 2009; Christopoulou et al. 2015). Only LsatNRC0 is not part of any of the reported MRCs (**Figure 2B**).

Due to a lack of chromosome-level assemblies for the wild lettuce species, we could not determine if the NRC-H and NRC-S in these genomes are genetically clustered together. However, our genome analysis revealed Ast-LvirNRC1 (Helixer_CAKMRJ010003334.1_001958.1) positioned near clade 2 and 3 sensors on contig2395 (**Figure 2A and S4**). We further examined five other species across Asterales to investigate whether NRC-S are in genetic clusters with the NRC helpers as a general trend (**Figure S5**). In *Cichorium intybus*, CiNRC0 (Helixer_CM042016.1_001204.1) is located within 500K base pair distance of a clade 1 NRC sensor on chromosome 8 (Helixer_CM042016.1_001201.1), whereas Ast-CiNRC1a (Helixer_CM042010.1_003195.1) and Ast-CiNRC1b (Helixer_CM042010.1_003196.1) are clustered with sensors from clade 2 and 3 on chromosome 3 (**Figure S6 and Data S6**). In *Cynara cardunculus*, CcNRC0 (Helixer_NC_037542.1_000084.1) is in a cluster with a clade 1 sensor (Helixer_NC_037542.1_001617.1) on chromosome 15, and Ast-CcNRC1 is in a genetic cluster with four clade 1 sensors on contig NW_020200580.1 (**Figure S7 and Data S6**). In *Helianthus annuus*, *Chrysanthemum lavandulifolium* and *Codonopsis lanceolata*, none of the helpers are clustered with any sensors (**Figures S6, S7 and Data S6**). These findings suggest variations in the physical clustering of NRC helpers and sensor clades within the *Lactuca* genus and between different Asterales species.

### LsatNRC0 and Ast-LsatNRC1 are required for cell death signaling of distinct NRC sensor phylogenetic clades

To investigate the functional connections of the lettuce NRC network and to test the phylogeny-based classification of these NLRs into NRC-S and NRC-H, we used Agrobacterium-mediated transient co-expression assays (Agroinfiltration) in an *nrc2/3/4* CRISPR knockout line of *Nicotiana benthamiana* (Wu et al. 2017). We agroinfiltrated autoactive mutants of the putative NRC-S and NRC-H to test their functionality, as the cognate effectors for these sensors remain unidentified. These mutants were generated by introducing a D-to-V mutation within the MHD motif of the NLRs, located near the C-terminus of the NB-ARC domain (Adachi et al. 2019a). We first tested whether transiently expressed NRC-S or NRC-H alone could induce cell death in *N. benthamiana*. While none of the NRC-S triggered cell death on their own, the autoactive mutants of the two lettuce NRC helpers displayed a macroscopic hypersensitive cell death response (**Figure S8A**). We then co-expressed the NRC-S and NRC-H in pairs to assess their potential functional connections and the dependence of lettuce NRC-S on the two NRC helpers. Among the 15 tested NRC-S, 9 displayed HR as autoactive mutants when co-expressed with one or both wild-type NRCs. Three NRC-S induced cell death through only LsatNRC0, while four responded solely through Ast-LsatNRC1. Notably, two sensors were functionally connected with both LsatNRC0 and Ast-LsatNRC1 (**Figure 3, S8B, Data S7**). While both NRC helpers possessed MADA motifs, none of the 15 tested sensors got a hit for MADA HMM profile (**Figure 3, Data S8**) (Adachi et al. 2019a). This was consistent with the cell death assays showing NRC helpers but not NRC sensors are able to cause cell death on their own.

**Figure 3.**
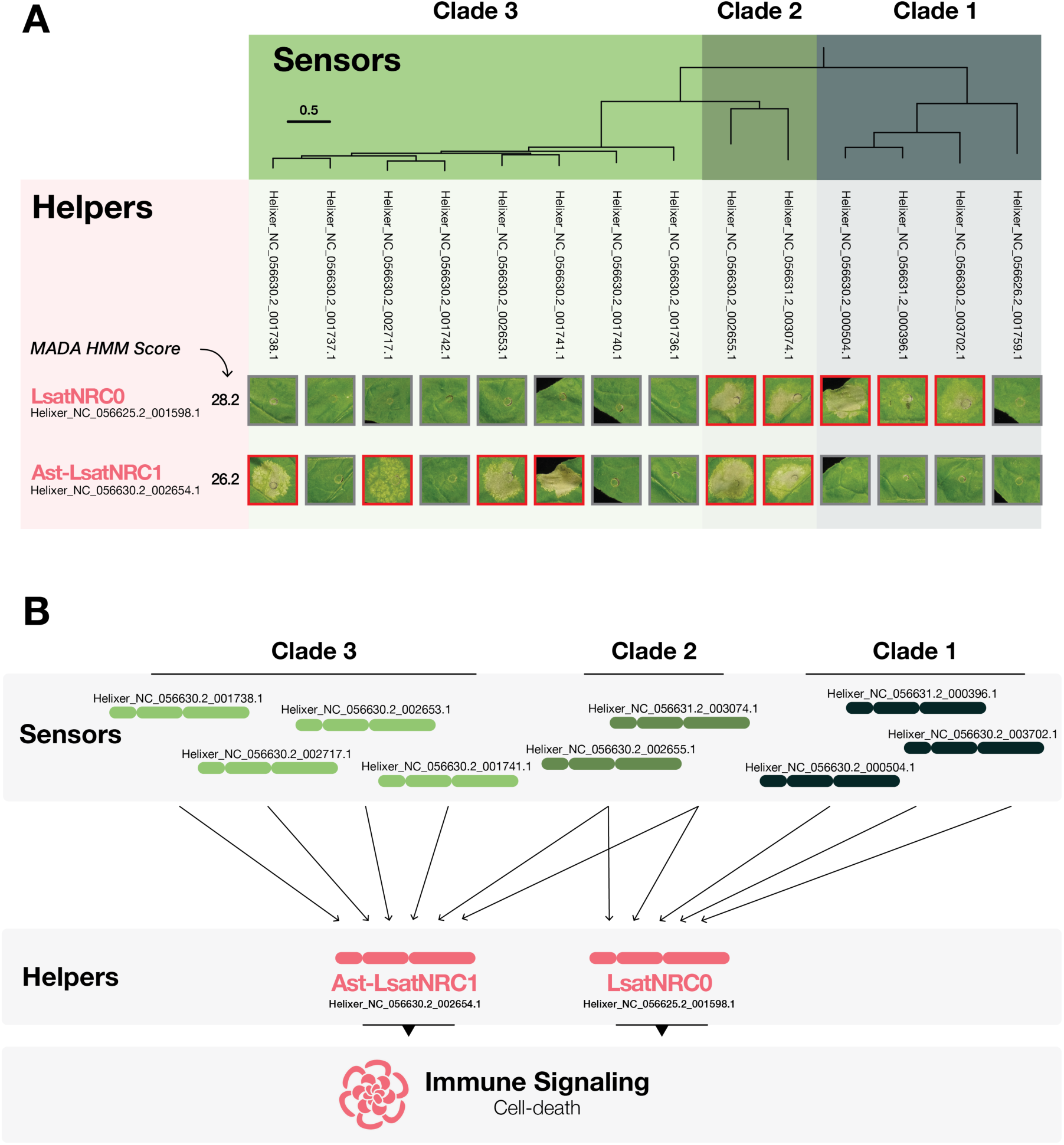
LsatNRC0 and Ast-LsatNRC1 mediate the cell death signaling of distinct NRC-S phylogenetic clades. (A) Co-agroinfiltration cell death assays of NRC-H and NRC-S. NRC-S expressed as autoactive mutants with wild-type NRC helpers. (B) Lettuce NRC network model. The 9 validated NRC-S can cause cell death through LsatNRC0 and Ast-LsatNRC1 in a partially redundant manner.

By analyzing the cell death phenotypes alongside the lettuce NRC network superclades’ phylogenetic tree, we observed a phylogenetic shift in helper dependency within the sensor clades. Specifically, all LsatNRC0-dependent sensors belonged to clade 1, while Ast-LsatNRC1-dependent sensors resided in clade 3. Interestingly, NRC-S that functioned with both helpers are clade 2 members positioned between clades 1 and 3 (**Figure 3A**). Taken together these observations indicate the presence of a functional NLR network with partially redundant signaling pathways in lettuce (**Figure 3B**).

### In contrast to NRC helpers, NRC sensors fail to form resistosome-like oligomers when modeled with AlphaFold 3

Recently, we demonstrated that AlphaFold 3 (Abramson et al. 2024) can differentiate between helper and sensor NLRs within pairs by predicting resistosome-like structures for helpers but not for sensors (Toghani et al. 2024). Building on these findings, we applied a similar approach to the lettuce NRC network superclade to investigate potential differences in the *in-silico* oligomerization of NRC-H and NRC-S. Using AlphaFold 3, we modeled five replicates of the two NRC helpers and 16 NRC-S sequences from lettuce as hexamers in the presence of 50 oleic acid molecules, simulating the plasma membrane environment. To evaluate these models, we extracted and analyzed key metrics, including pLDDT (per-residue measure of local confidence), pTM (predicted template modeling score for overall structure), ipTM (interface-specific pTM), and per-chain pTM (**Data S8**). AlphaFold 3 successfully modeled resistosome-like oligomers for both NRC helper sequences, though it was unable to generate a complete N-terminal funnel in any of the LsatNRC0 predicted structures (**Figure 4**).

**Figure 4.**
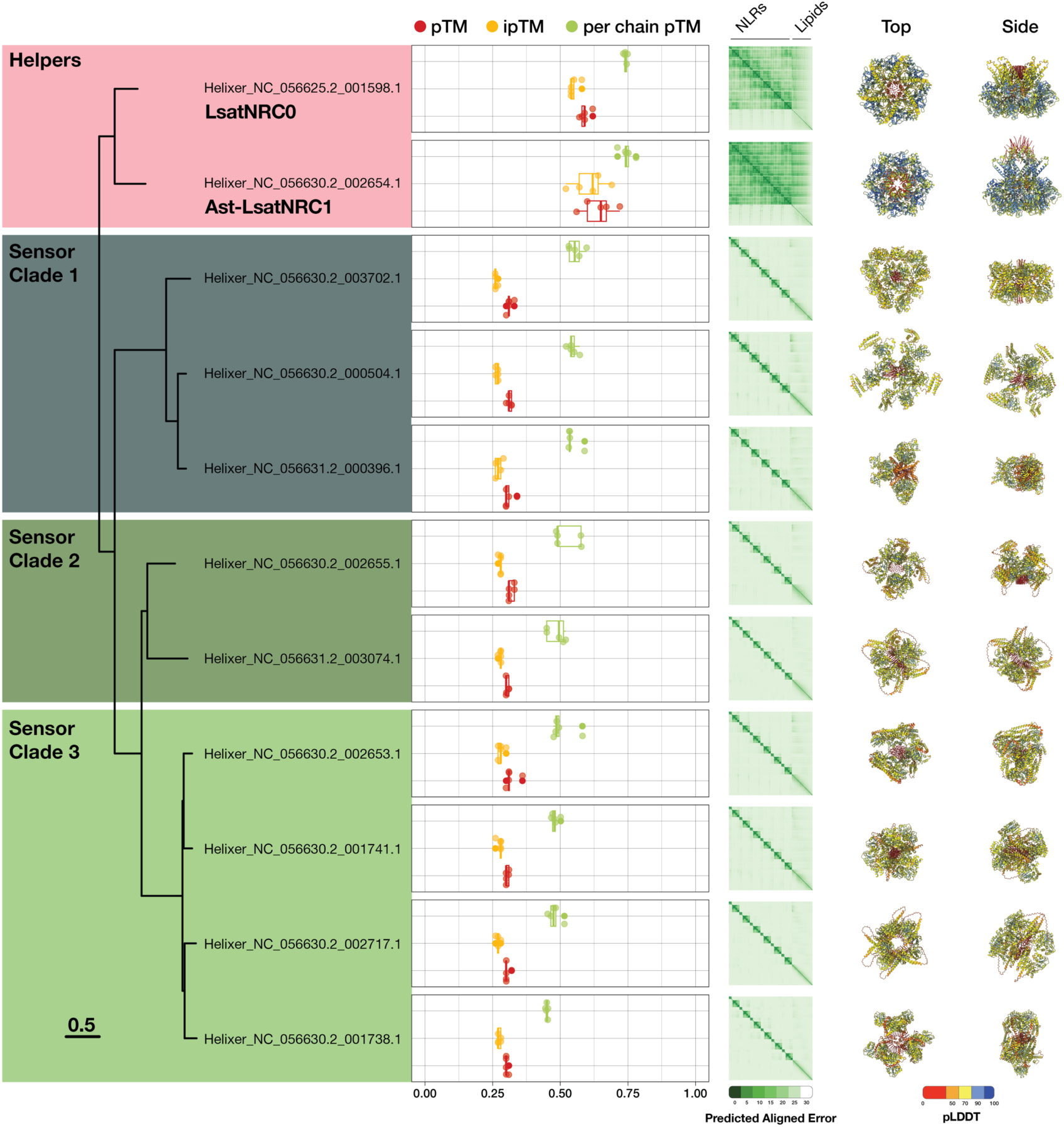
In contrast to NRC helpers, NRC-S failed to form resistosome-like structures when modeled with AlphaFold 3. Structural modeling with AlphaFold 3 was used to predict the ability of the tested lettuce NRC-H and NRC-S to form resistosome-like hexameric oligomers. Simulations included 50 oleic acid molecules to approximate the plasma membrane. Key structural metrics, including pLDDT (per-residue measure of local confidence), pTM (predicted template modeling score for overall structure), ipTM (interface-specific pTM), and per-chain pTM, are plotted for each protein (n=5). Example predicted structures from the fourth replicate are displayed in top and side views, colored with the pLDDT color scale indicating local confidence levels.

This was consistent with our previous observations when modeling NRC0 oligomers (Madhuprakash et al. 2024). In contrast, none of the NRC-S from the three sensor subclades formed resistosome-like structures (**Figure 4**). Across all replicates, NRC-S models consistently showed low confidence with high predicted aligned errors (**Figure 4 and S9**). Structurally, NRC-S displayed lower metrics compared to NRC helpers, with pTM and ipTM values below 0.5 and per-chain pTM values below 0.7 (**Figure 4, S9**). Conversely, helpers had consistently higher metrics, with pTM and ipTM exceeding 0.5 in all replicates and an average per-chain pTM of 0.75 (**Figure 4, S9**). These results suggest that AlphaFold 3 can distinguish between NRC-S and NRC-H based solely on sequence and their capacity to form higher-order resistosome-like oligomers.

### Highly variable sites localize to different domains in different NRC-S subclades

Despite having a common evolutionary origin, NRC sensors and helpers exhibit functional diversity (Adachi et al. 2019b; Contreras et al. 2023a). This prompted us to investigate the sequence diversification of NRC-H and NRC-S in the *Lactuca* genus. To test this, we performed Shannon’s entropy analysis on each of the helper and sensor clades (Hausser and Strimmer 2009). Shannon’s entropy measures the variability of amino acids at a specific position in a protein sequence alignment. A value of 0 indicates only one amino acid is present, while a value of 4.32 signifies all 20 amino acids are present with equal frequency. We used entropy values of 1.5 and higher as a threshold for identifying highly variable residues as reported in previous NLR studies (Prigozhin and Krasileva 2021; Selvaraj et al. 2024). The *Lactuca* NRC helper clade exhibited minimal variation across its entire length, with only 23 residues exceeding the 1.5 entropy threshold. These variable residues were primarily located within the CC domain. Conserved motifs such as MADA, P-loop, and MHD also had low entropy values (**Figure 5A and 5B, Data S10**). In contrast, the sensor clades displayed significant variations: NRC-S clades 1 and 3 contained a substantially higher number of variable residues (146 and 128, respectively) compared to clade 2 (28 residues) that surpassed the 1.5 entropy threshold. Within clade 1, three clusters of highly variable residues were observed: one in the middle of the CC domain, and two near the N-termini of the NB-ARC and LRR domains, respectively (**Figure 5A and 5B, Data S10**). These findings corroborate the previous report on the presence of variable regions in these domains of NRC0-dependent sensors (Sakai et al. 2024). In clade 2, the few variable residues are scattered throughout the NLR sequence. Interestingly, clade 3 displayed a concentration of variable residues within the concave surface of the LRR domain repeats (**Figure 5A and 5B, Data S10**). As mentioned previously, we did not detect the MADA motif in NRC-S using the MADA HMM. However, despite differences in diversification patterns, all sensors have a conserved “<u>MAYAxVQMFMEKLKQLIYx</u>” N-terminal motif resembling the NRC0-S N-terminal motif reported previously (**Figure 5A**) (Sakai et al. 2024). P-loop and MHD motifs are also highly conserved between all NRC-S (**Figure 5A**). Together these indicated not only helpers and sensors exhibit different patterns of diversification but also NRC-S subclades, compared to each other, have diversified and evolved differently.

**Figure 5.**
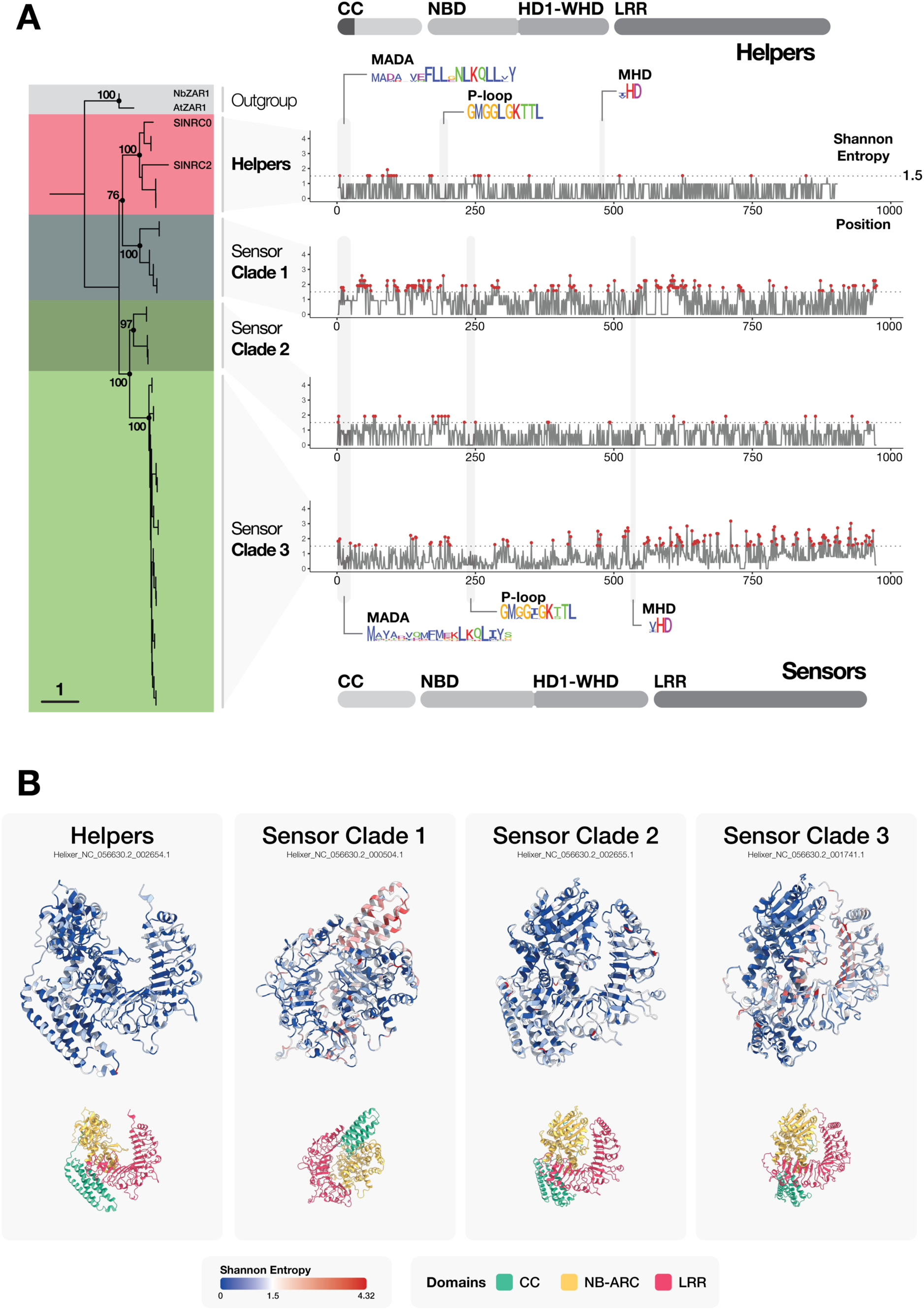
Lettuce NRC helpers and sensors have differential patterns of diversification. (A) Shannon entropy plots of the full-length NLR sequences calculated within each NRC subclade. Highly variable residues with Shannon entropy values of 1.5 and higher are highlighted by red dots. (B) Predicted AlphaFold 3 structures of representative sequences from each NRC subclade colored by Shannon entropy values (top) and domains (bottom).

### Sensors dependent on Ast-LsatNRC1 are under the higher positive selection compared to LsatNRC0-dependent sensors

To examine the selection pressures that affected mutations in the NRC helper and NRC-S subclades, we conducted positive selection tests using CodeML in PAML v4.10.7 software (Yang 1997). Six models were evaluated for each NRC subclade (helpers, sensors C1, C2, and C3): M0 (one-ratio), M1 (nearly neutral), M2 (positive selection), M3 (discrete), M7 (beta), and M8 (beta + ω). Likelihood ratio tests (LRT; 2ΔL = 2(InL_alternative_ hypothesis – InL_null hypothesis_)) compared M2 vs M1, M3 vs M0, and M8 vs M7 for each clade (**Data S11**). The LRT values were evaluated against the *χ*2 critical thresholds to assess significance. A higher LRT than the critical value indicates that the alternative hypothesis better fit the data.

For the helper clade, the calculated LRT for M2 vs M1 (0) and M8 vs M7 (6.379) did not exceed the *χ*2 critical threshold (9.21 at 1% significance; degrees of freedom = 2; **Data S11**). Conversely, the LRT for M3 vs M0 (69.097) surpassed the threshold (13.28 at 1% significance; degrees of freedom = 4; **Data S11**), demonstrating that the M3 (discrete) model fit the data significantly better than M0 (neutral). The M0 model inherently excludes sites under diversifying selection (*ω* > 1). However, no sites under positive selection were identified in the alternative models (M2, M3, M8) (**Figure 6A; Data S11**).

**Figure 6.**
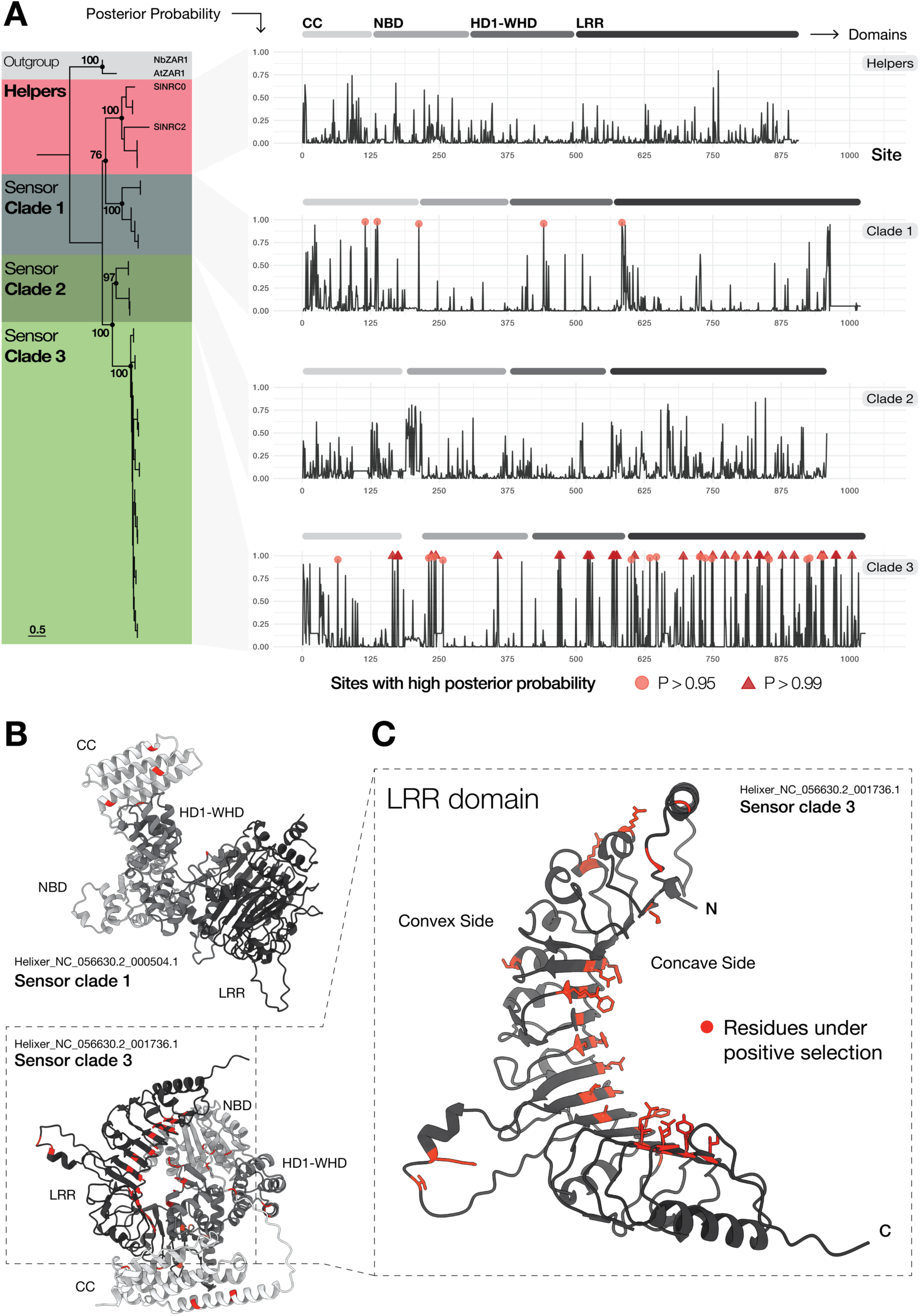
Ast-LsatNRC1-dependent sensors are under more positive selection compared to NRC0-dependent sensors and NRC helpers. (A) Posterior probabilities for full-length NLRs in each NRC clade, estimated using the M8 (beta + *ω*) model from the CodeML package in PAML. Sites with posterior probabilities (P) > 0.95 and > 0.99 are marked by circles and triangles, respectively (Yang 1997). (B) Sites under positive selection (red) mapped onto predicted structures of representative sensor NLRs from clades 1 and 3, generated using AlphaFold 3. (C) Zoomed-in view of the LRR domain from a representative clade 3 sensor NLR, highlighting sites under positive selection (red).

For sensor clade 1, all three LRT comparisons yielded values exceeding the *χ*2 thresholds, supporting the alternative models as better fits to the data (M2 vs M1 = 22.079, M3 vs M0 = 392.939, M8 vs M7 = 58.949; **Data S11**). Bayes Empirical Bayes (BEB) analysis (Yang et al. 2005) under the M8 (beta + *ω*) model revealed five sites under positive selection (posterior probability > 0.95). Mapping these to an AlphaFold 3-predicted structure of a representative clade 1 sensor NLR (Helixer_NC_056630.2_000504.1) localized three of these sites to the CC domain (**Figure 6B**).

In sensor clade 2, the LRT for M2 vs M1 (6.806) was below the *χ*2 critical value. In contrast, LRTs for M3 vs M0 (21.639) and M8 vs M7 (157.960) exceeded the critical thresholds, favoring the alternative models (**Data S11**). Despite this, BEB analysis under the M8 model did not identify any sites under positive selection (posterior probability > 0.95) (**Figure 6A; Data S11**).

For sensor clade 3, all three LRT comparisons supported the alternative models (M2 vs M1 = 337.837, M3 vs M0 = 1372.561, M8 vs M7 = 400.550; **Data S10**). BEB analysis from the M8 model identified 48 sites under positive selection (posterior probability > 0.95) (**Figure 6A; Data S11**). Mapping these onto an AlphaFold 3-predicted structure of a representative clade 3 sensor NLR (Helixer_NC_056630.2_001736.1) revealed that 33 of the 48 sites were located within the LRR domain (**Figure 6C**). Notably, most of these sites clustered on the concave face of the LRR, a region implicated in effector binding based on previous studies identifying signatures of positive selection and high variability (**Figure 6C**) (Michelmore and Meyers 1998; Prigozhin and Krasileva 2021).

Next, we estimated the pairwise nonsynonymous (*d*_N_) and synonymous (*d*_S_) substitution rates and *d*_N_/*d*_S_ ratios (*ω*) for members of each NRC subclade using the Nei and Gojobori (1986) method (**Data S12**) (Nei and Gojobori 1986; Yang and Nielsen 2000). Analyses were conducted on full-length sequences and individual domains (CC, NB-ARC, and LRR), where *ω* > 1 implied presence of positive selection. Among full-length and domain-based comparisons, NRC helpers had lower average ω compared to all NRC-S clades (**Figure S11**). While sensor clade 3 on average had overall higher ω than the rest of NRC-H and NRC-S clades, the CC domain in clade 1 sensors had *ω* > 1 values, indicative of positive selection (**Figure S11**). Overall, the pairwise *d*_N_/*d*_S_ ratios of NRC-H and NRC-S subclades further supported the diversification and positive selection patterns observed in the entropy and CodeML positive selection analyses, respectively.

## DISCUSSION

The NRC network originated approximately 100 million years ago within the superasterid lineage of land plants (Wu et al. 2017). In this study, we conducted a comparative phylogenetic and diversification analysis of the NRC network superclade in the Asterales and Solanales orders of the Asterid clade. For this purpose, we used a *de novo* generated annotation dataset for 40 Solanales and 29 Asterales genomes using the deep-learning-based Helixer software, ensuring comparable and harmonized gene models across all analyzed genomes (Stiehler et al. 2021; Holst et al. 2023). We then focused on common lettuce as a model Asterales species and experimentally validated a functional and partially redundant NRC network that was identified using phylogenetics analysis. We used this minimal NRC network to study the diversification and evolution of different NRC-H and NRC-S subclades.

The NRC network has expanded significantly in the lamiid lineage of Asterids over tens of millions of years, whereas the campanuliid lineage, including Asterales, has experienced much less expansion (Goh et al. 2024). Our analysis of 40 Solanales and 29 Asterales genomes revealed marked differences in NRC network evolution between these orders. (**Figure 1**). Notably, NRC superclade NLRs make up only 6.6% of the total NLR repertoire in Asterales, compared to 50.3% in Solanales (**Figure S2**). Despite similar average total NLR counts (338.6 in Solanales vs. 300 in Asterales), Asterales appears to have expanded other NLR clades, such as TIR-NLRs. In lettuce, for example, TIR-NLRs comprise a majority of the NLRome (57.4%; 206/359), while NRC superclade sequences represent only 5% (18/359) (**Figure S2 and S3**). This aligns with previous findings of extensive proliferation of NRC in lamiids versus campanuliids (Goh et al. 2024). Our study found no evidence of Rx-type sensors in Asterales, a class significantly expanded in Solanales and linked to many characterized R-genes in Solanaceous species (Wu et al. 2017).

Although Asterales SD-type sensors phylogenetically cluster with their Solanales counterparts, they lack the N-terminal SD-domain. It was previously suggested this is due to a loss of the sensors with SD integration in Asterales (Seong et al. 2020). However, given the presence of SD-type-like sensors in sugarbeet (*Beta vulgaris*), a Caryophyllales species, it’s more likely that phylogenetically close sensors, with and without the SD integration, were already present in the common ancestor of superasterids (Seong et al. 2020), but the sensor with the SD integration was possibly lost in the ancestral Asterales species (**Figure 1**). More in-depth phylogenetic analysis across superasterids may provide insights into the origin of SD-type sensors and possible SD integration/emergence events.

In lettuce, NRCs are divided into four distinct phylogenetic subclades: a helper clade and three sensor clades (C1, C2, and C3) (**Figure 2**). Mapping these phylogenetic clades to the classification by Meyers et al. (1998) revealed their alignment with four subclades (RGC7, RGC9, RGC26, and RGC27) (Meyers et al. 1998). Notably, Ast-LsatNRC1 (RGC7), all clade 3 sensors (RGC9), and a single clade 2 sensor (RGC27) are part of MRC8c, which has been linked with QTLs for resistance to *Fusarium oxysporum* and *Verticillium dahliae* (Christopoulou et al. 2015). Other NRC-S from clades 1 and 2 are associated with Major Resistance Clusters (MRC) 4, 8a, and 9a, while *NRC0* is not linked to any known MRC. Although MRC4 and 9a have been implicated in downy mildew resistance, to date, NRC superclade NLRs have not yet been implicated in resistance to pathogens in common lettuce (Meyers et al. 1998; Christopoulou et al. 2015). However, further investigation into the role of NRC network components classified as MRC8c in resistance against *F. oxysporum* and *V. dahliae* would be valuable, as they represent 12 out of 26 immune receptor genes in this genetic cluster (Christopoulou et al. 2015).

*NRC0* is the most conserved NRC helper across superasterids (Goh et al. 2024; Sakai et al. 2024), while other NRC helpers, such as NRC2, NRC3, and NRC4 in Solanaceae, show lineage-specific expansions (Goh et al. 2024; Selvaraj et al. 2024). Although Asterales have a less extensive overall expansion of NRCs compared to Solanales, many species in Asterales retain an additional NRC helper besides *NRC0*. Previous studies have demonstrated that NRC helpers, including *NRC0* and *NRC6*, can be genetically clustered with NRC sensors in several plant species (Lüdke et al. 2023; Sakai et al. 2024). However, in common lettuce, while Ast-LsatNRC1 is clustered with clade 2 and 3 sensors, *LsatNRC0* is not clustered with any sensor NLRs (**Figure 2B**). Further examination across Asterid species revealed species-specific patterns in the genetic linkage of NRC superclade NLRs. For instance, in *Cichorium intybus* (chicory), *CiNRC0* is proximal to a clade 1 sensor, and both *Ast-CiNRC1* copies cluster with clade 2 and 3 sensors. However, in *Helianthus annuus* (sunflower), no helpers are closely associated with sensors (**Figure S6**). These observations suggest that genetic linkage of *NRC0* with sensors is ancestral (Sakai et al. 2024), but the physical clustering of NRC superclade NLRs may be influenced by species-specific genomic rearrangements.

We expanded Goh et al. (2024) analyses by determining the functional dependency of lettuce NRC sensors on NRC helpers using a transient co-expression system in *N. benthamiana* (Goh et al. 2024). Of the 15 NRC-S tested, 9 triggered hypersensitive cell death when co-expressed with at least one lettuce NRC helper (**Figure 3 and S8**). Phylogenetic analysis revealed a shift in helper dependency among lettuce sensors: Clade 1 sensors exclusively rely on LsatNRC0, while clade 3 sensors depend solely on Ast-LsatNRC1. Clade 2 sensors, an intermediate group between clades 1 and 3, can signal through both LsatNRC0 and Ast-LsatNRC1 (**Figure 3**). This gradual evolutionary shift in helper dependency provides us a simplified system to study the diversification and function of different NRC clades.

Previously, we demonstrated that in monocot NLR pairs, helpers could form resistosome-like oligomers when modeled using AlphaFold 3, whereas sensor NLRs were unable to form similar structures (Toghani et al. 2024). However, whether this finding extends to dicots and NLR networks remained unclear. In this study, none of the tested NRC-S triggered cell death when expressed on its own (**Figure S8**).

Multiple studies have also shown that NRC-S cannot independently induce cell death or form higher-order complexes before or after activation (Contreras et al. 2022; Ahn et al. 2023; Selvaraj et al. 2024). Given these observations, we modeled NRC-H and NRC-S as hexameric complexes using AlphaFold 3. Similar to results with sensor-helper pairs in rice, NRC helpers were predicted to form resistosomes, whereas NRC-S failed to yield meaningful or high-confidence structural predictions (**Figure 4**). Combined with our functional data and biochemical evidence from other NRC-S, this finding is consistent with the view that NRC-dependent sensors have lost the ability to form resistosomes (Contreras et al. 2022; Ahn et al. 2023). However, further structural studies on NRC-S in their resting and active states are needed to elucidate the mechanisms underlying sensor activation and sensor-helper interactions.

Our Shannon entropy and selection analyses revealed that NRC helpers generally show low variability and lack any signatures of positive selection, consistent with helper NLRs like ADR1, which are classified among low-variability NLR clades (**Figures 5, 6, S10, and S11**) (Seong et al. 2020; Prigozhin and Krasileva 2021; Sutherland et al. 2024). In contrast, NRC-S clades demonstrate clade-specific variation patterns. Clade 1 sensors, reliant on NRC0, show high variability in the CC domain and the N-terminal part of the LRR domain, aligning with previous findings by Sakai et al. (2024) (**Figures 5, 6, and S11**) (Sakai et al. 2024). The cause of this diversification—whether due to relaxed selection or co-evolution with pathogens—remains unclear. Clade 2 sensors exhibit generally low variation and can signal through both LsatNRC0 and Ast-LsatNRC1. Clade 3 sensors, dependent on Ast-LsatNRC1, display highly variable regions concentrated on the concave surface of the LRR domain, with numerous sites under positive selection, resembling highly variable NLRs like RPP1 in *Arabidopsis thaliana* (**Figures 5, 6, and S11**) (Prigozhin and Krasileva 2021). The observed genetic clustering within clade 3 suggests that recent tandem duplication events and accelerated co-evolution with pathogens may have driven their diversification (Barragan and Weigel 2021).

Why have NRC network clades evolved differently? The differential evolutionary trajectories of NRC network clades are shaped by their distinct roles, sensor-helper dependencies, and interactions with pathogens. The contrasting conservation of NRC0 and its functionally connected sensors versus the diversification of other NRC-H and NRC-S highlights varying selective pressures acting on these groups. NRC0 and its dependent sensors are conserved across Asterid lineages (Goh et al. 2024; Sakai et al. 2024) (**Figure 7**). The limited copy number and conservation of NRC0-dependent sensors, particularly in their LRR domains (which usually mediate pathogen recognition), may reflect detection of conserved effector molecules or guarding essential host components that are targeted by effectors, analogous to the ZAR1-ZRK system (Adachi et al. 2023). This conservation may limit the diversification of corresponding plant receptors. Such constraints could also arise from the need to maintain compatibility with NRC0, ensuring that evolutionary changes do not disrupt functional interactions or downstream signaling processes (**Figure 7**). In contrast, other NRC helpers and their associated sensors, especially those in lineages like Solanales, show greater diversification and expansion. This may stem from selective pressures such as detection a broader range of pathogen effectors, evading suppression, or introducing redundancy in signaling pathways (**Figure 7**) (Kamoun 2018; Adachi et al. 2019b; Derevnina et al. 2021; Contreras et al. 2023b; Sugihara et al. 2024). Non-NRC0-dependent sensors, often clustered together, exhibit strong signatures of positive selection, particularly in their LRR domains (**Figure 6C**), which likely reflects their engagement in a faster co-evolutionary arms race with pathogens compared to NRC0-dependent sensors.

**Figure 7.**
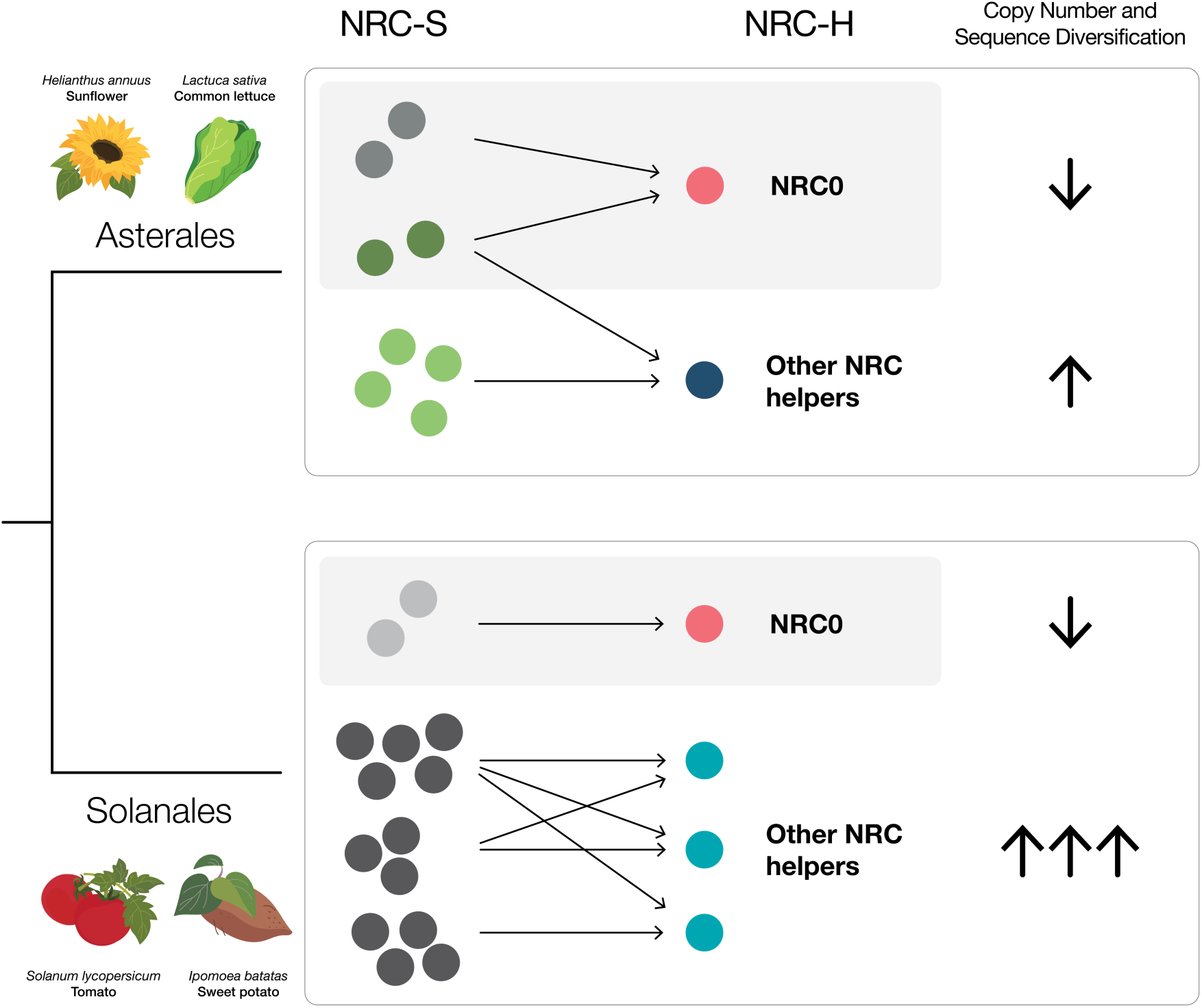
Evolutionary constraints and diversification patterns in NRC helpers and sensors. Sensors dependent on NRC0 are characterized by lower copy number and are under lower positive selection, likely due to evolutionary constraints associated with their dependency on NRC0. In contrast, other NRC helpers and the sensors depending on them have undergone significant expansions, exhibiting higher copy numbers and diversification rates. Representative species from each order are shown.

In summary, our study highlights the contrasting evolutionary trajectories between helper and sensor NLRs, as well as among distinct sensor NLR clades within the NRC network. The NRC network mediates robust immune responses against a diverse array of pathogens, including bacteria, fungi, oomycetes, nematodes, and viruses. Certain nodes, such as *NRC6*, demonstrate specialization by providing tissue-specific or pathogen-specific resistance (Lüdke et al. 2023). This multi-paced evolution of helpers and sensor clades has likely granted the network the flexibility to diversify and counter a broad spectrum of pathogens while allowing specialization and potential adoption of novel immune functions. The simplified Asterales NRC network in lettuce provided insights into how a shift in helper dependency may have facilitated sensor diversification and expansion distinct from their more conserved ancestral counterparts (**Figure 7**). Furthermore, our findings illustrate how closely related NLRs with a functional connection, such as lettuce NRCs, can exhibit varying diversification patterns, similar to those seen in more distantly related clades or in NLRs displaying allelic diversity within a single locus, such as RGC2 (Kuang et al. 2004). Addressing the molecular mechanisms governing helper specificity and sensor-helper interactions in future studies will illuminate how these immune receptors have adapted and evolved across plant lineages. This will facilitate breeding and immune engineering efforts in Asterid crop species.

## METHODS

### Extraction of NRC network superclade

A total number of 29,261 NLRs were compiled from the NLRtracker output of 39 Solanales and 29 Asterales *de-novo* annotated high-quality genome assemblies from NCBI (https://www.ncbi.nlm.nih.gov/datasets/) and a *S. melongena* chromosome-level genome assembly from GWH genome warehouse datasets (Chen et al. 2021) (**Table S1**) (Toghani et al. 2025a, 2025b). Based on the simple domain annotation from NLRtracker we only kept NLRs with “CNL”, “CNLO”, “CN”, “OCNL”, “CONL”, “NL”, “NLO”, “ONL”, “BCNL”, “BNL”, “BCN”, “BCCNL”, “BNLO”, “BOCNL”, “RNL”, “TN”, “TNL”, “TNLO”, “TNLJ” domain architectures. This led to keeping 25,042 NLRs. The remaining NLRs were then deduplicated to keep the 24,518 non-redundant sequences. Finally, based on NLRtracker domain output, any entry with NB-ARC domain shorter than 250 amino acids and longer than 400 amino acids were removed. This led to a final 21,232 sequences that was used for the phylogenetics analysis.

To construct a phylogenetic tree of the extracted NLRs, the NB-ARC domain sequences were aligned with RefPlantNLR NB-ARCs using FAMSA v2.2.2 [default options] (Deorowicz et al. 2016). The alignment was then used to construct a phylogenetic tree using FastTree v2.1.11 [-lg] (Price et al. 2010; Kourelis et al. 2021). The NRC network superclade along with CcRPP1 paraphyletic clade, including 7,040 sequences, was extracted based on the presence of reference sequences from a well-supported branch containing NRC-H and NRC-S. The NB-ARC domain of the extracted sequences were aligned using MAFFT v7.525 (Katoh and Standley 2013). The alignment was used to construct a new phylogenetic tree in the same way as before. The resulting tree was rooted at the CcRPP1 clade.

### Phylogenetics and gene distance analyses

Plant taxonomy trees and taxa divergence times were obtained from TimeTree 5 database (timetree.org) (Kumar et al. 2022). 122 NRC sequences including helpers and sensors from *Lactuca sativa*, *Lactuca saligna*, *Lactuca virosa*, *Cichorium intybus*, *Helianthus annuus*, *Chrysanthemum lavandulifolium*, *Cynara cardunculus*, and *Codonopsis lanceolata* present in the dataset were extracted based on phylogeny. The NB-ARC sequences of extracted proteins were aligned with AtZAR1, NbZAR1, NbNRC2, SlNRC0, as references, using MAFFT v7.525 [--anysymbol] (Katoh and Standley 2013). FastTree v2.1.11 [-lg] was used to generate the tree of NRC network in the mentioned species (Price et al. 2010). We then used IQtree v2.3.0 [-B 1000 -m MFP] to generate a phylogenetic tree of 40 sequences in the NRC network in the *Lactuca* genus (Kalyaanamoorthy et al. 2017; Minh et al. 2020). The resulting tree was divided to helpers, sensor clade 1, sensor clade 2, and sensor clade 3 subclades based on branch lengths and reference sequences.

To obtain the physical location of the genes on the chromosome, first the GFF output from Helixer was simplified using GffRead [default options] and then imported into R using rtracklayer package (Lawrence et al. 2009; Pertea and Pertea 2020). The physical distances between genes was calculated using a custom R script and gene coordinates were exported to be visualized on chromosomes using MG2C online tool (Chao et al. 2021). The output was then visualized manually.

### Retrieving previous lettuce NLRome classifications

Lettuce NLRome IDs from Christopoulou, et al., 2015, and Lettuce V8 genome assembly from Phytozome 13 were extracted (Goodstein et al. 2012; Christopoulou et al. 2015; Reyes-Chin-Wo et al. 2017). We then used BLASTP as part of BLAST+ v2.16.0 to identify corresponding gene identifiers in the Phytozome 13 annotation to the new Helixer annotation (Camacho et al. 2009). The BLAST output was then processed in R and the gene coordinates, major resistance loci information, and RGC groups were retrieved for the new Helixer annotation.

### Agroinfiltration and cell death assays

To test the functionality of lettuce NRCs, annotated genes were cloned using the Golden Gate Modular Cloning (MoClo) kit (Weber et al. 2011) and the MoClo Plant Parts kit (Engler et al. 2014). In brief, coding sequences were synthesized by GENEWIZ as Golden Gate Level 0 pICH41155 modules and subsequently transferred to the binary vector pICH86988 through *Bsa*I digestion (Weber et al., 2011). Native *Bsa*I sites in the original sequences were domesticated as needed. In addition to wild-type clones, a D-to-V mutation within the MHD motif was introduced to generate autoactive versions (Adachi et al. 2019a) of the lettuce NRCs. Verified clones were transformed into *A. tumefaciens* strain GV3101 pMP90 for ectopic expression in *N. benthamiana*.

For cell death assays, wild-type and nrc2/3/4 *N. benthamiana* plants were grown in a controlled environment growth chamber at 22–25°C with a 16-hour light period and 45%/65% relative humidity during the light/dark cycle. Light intensity was approximately 200 µmol/m^2^/s. Four-to five-week-old plants were used for agroinfiltration, following the methods described by Bos et al. (2006) (Bos et al. 2006). The final OD_600_ of all Agrobacterium suspensions was adjusted to 0.5 in infiltration buffer (10mM MES, 10mM MgCl_2_, and 150µM acetosyringone, pH5.6). This concentration was used for both expressing clones individually and co-expressing NRC-H and NRC-S of interest. The cell death phenotype was imaged 4 to 5 days post-agroinfiltration.

### Sequence polymorphism and selection analyses

To calculated the Shannon entropy values for each clade, we used the “entropy” R package as part of a custom script as described before (Hausser and Strimmer 2009; Selvaraj et al. 2024). For this purpose, the helper subclade, and all sensor clades from *Lactuca* genus were aligned separately using MAFFT v7.525 [--anysymbol--localpair]. The sensor clade alignment was trimmed with ClipKit v2.0.1 [-m gappy] as described before and the sensor subclades were extracted from the resulting alignment. The entropy values for each clade were then visualized using a ggplot2 (Wickham 2016). Sequence logos for known conserved motifs in helper and sensor clades were generated using ggseqlogo R package (Wagih 2017).

Nucleotide sequence of the NRC superclade sequences were obtained using a custom python script from a compiled CDS file of the annotated genomes. Codon-based nucleotide alignments were generated for each subclade generated using MACSE v2.07 [-prog alignSequences-seq] (Ranwez et al. 2018). Positive selection tests were conducted on resulting alignments using CodeML in PAML v4.10.7 (Yang 1997). Six models were applied for each subclade: M0 (one-ratio), M1 (nearly neutral), M2 (positive selection), M3 (discrete), M7 (beta), and M8 (beta + ω). Likelihood ratio tests (LRT; 2ΔL = 2 (InL_alternative_ hypothesis – InL_null hypothesis_)) were performed to compare M2 vs. M1, M3 vs. M0, and M8 vs. M7 for each clade. LRT values were evaluated against χ2 critical thresholds to assess significance, with higher LRT scores indicating a better fit for the alternative hypothesis. Bayes Empirical Bayes (BEB) analysis under the M8 model, included in the CodeML output, identified sites under positive selection with probabilities higher than 0.95 (Yang et al. 2005).

Pairwise nonsynonymous (*d*_N_) and synonymous (*d*_S_) substitution rates were estimated for full-length sequences and individual domains (CC, NB-ARC, and LRR) in each clade using the Nei and Gojobori (1986) method (Nei and Gojobori 1986; Yang and Nielsen 2000). These rates were extracted from the “2NG.dN” and “2NG.dS” outputs of CodeML using a custom R script. All figures were generated with the ggplot2 package in R (Wickham 2016).

### Protein structure prediction and mapping Shannon entropy to protein structures

AlphaFold 3 [Seed=1, Ligand=ADP] was used to predict the monomeric structures of representative sequences from each clade (**Data S9**) (Abramson et al. 2024). The Shannon entropy values for each NRC subclade was mapped to a representative sequence from each clade using a custom R script and ChimeraX (Pettersen et al. 2021). For oligomeric structures, AlphaFold 3 with [Ligand=OLA x 50] was used to model sequences (**Data S9**). First model for each sequence was modeled with Seed = 1. Four other replicates were modeled for each sequence with random seeds. Model confidence and metadata were processed and plotted with R. All scripts available at [https://github.com/amiralito/lettuce_salad].

## DATA AVAILABILITY

Additional supplementary files and material are available at Zenodo repository (https://doi.org/10.5281/zenodo.14544899) (Toghani and Kamoun 2025). Genome annotations and sequence data available at Dryad repository (https://doi.org/10.5061/dryad.sxksn03d6; https://doi.org/10.5281/zenodo.14720919) (Toghani et al. 2025b, 2025a).

## ACKNOWLEDGMENTS

We thank Daniel Lüdke, Joe Win, Jonathan D. G. Jones (The Sainsbury Laboratory, UK), Philip Carella (John Innes Centre, UK), and Chandler Sutherland (University of California, Berkeley, USA) for their valuable and technical feedback. A.T. thanks Josh W. Bennett (John Innes Centre, UK) for insightful scientific discussions and C. Blasco for precision and detailed feedback. We also deeply appreciate the invaluable support and contributions of the TSL community and support groups.

## AUTHOR CONTRIBUTIONS

Conceptualization: A.T., S.K., H.A., R.F.

Data Curation: H.P., A.T., T.S., H.A.

Formal Analysis: H.P., A.T., T.S., H.A.

Investigation: H.P., A.T., T.S., H.A.

Methodology: H.P., A.T., A.P., T.S., H.A., Y.S., J.K

Resources: A.T., Y.S., S.K., H.A.

Software: A.T., A.P., T.S., H.A., Y.S.

Supervision: A.T., S.K.

Funding Acquisition: S.K.

Project Administration: A.T., S.K.

Writing Initial Draft: A.T., H.P.

Editing: A.T., S.K., H.A., Y.S., M.P.C

## SUPPLEMENTARY MATERIAL

**Data S1.** List of genome assemblies and species used in this study.

**Data S2.** List of NLR sequences and metadata used for the phylogenetic analysis.

**Data S3.** List of NRC helper and sensor sequences used for the phylogenetic analysis.

**Data S4.** List of *Lactuca* NRC helper and sensor sequences and metadata.

**Data S5.** List of selected Asterales NRC helper and sensor sequences and metadata.

**Data S6.** Gene-distance matrix of NRC helpers and sensors.

**Data S7.** Raw cell death scores of tested lettuce NRC helpers and sensors in *Nicotiana benthamiana*.

**Data S8.** HMMsearch output of MADA motif for NRC superclade sequences in *Lactuca* genus.

**Data S9.** Model confidences for the predicted models used in different analysis.

**Data S10.** Entropy values for NRC subclades.

**Data S11.** PAML results tables for the NRC subclades.

**Data S12.** Pairwise *d*_N_ and *d*_S_ values for the four NRC subclades.

---

**Figure S1.**
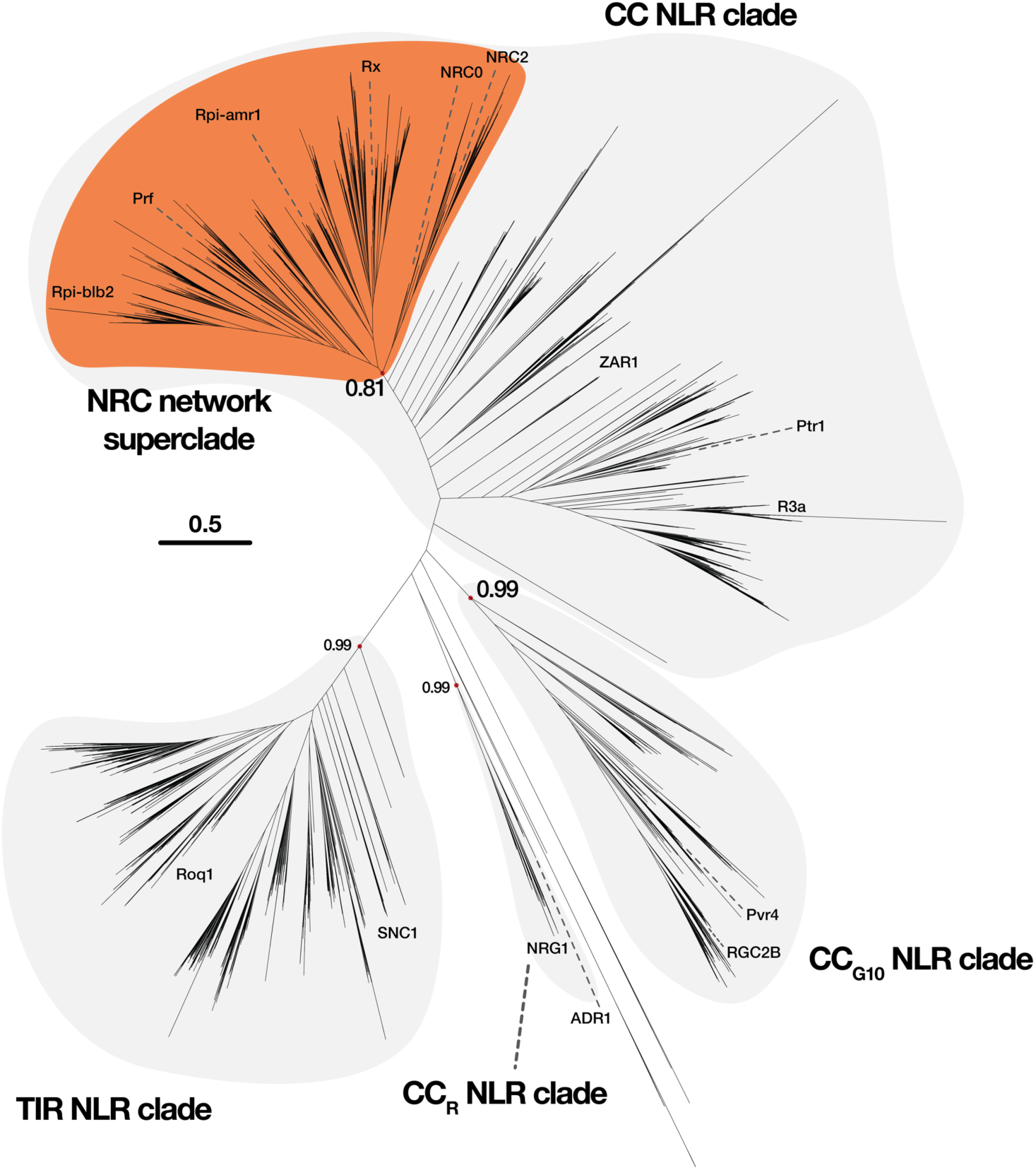
Phylogenetics tree of 21,645 Solanales and Asterales NLRs with RefPlantNLR as the reference. NRC network superclade, consisting of NRC helpers (NRC-H) and sensors (NRC-S), forms a well-supported clade within the CC-NLR clade.

**Figure S2.**
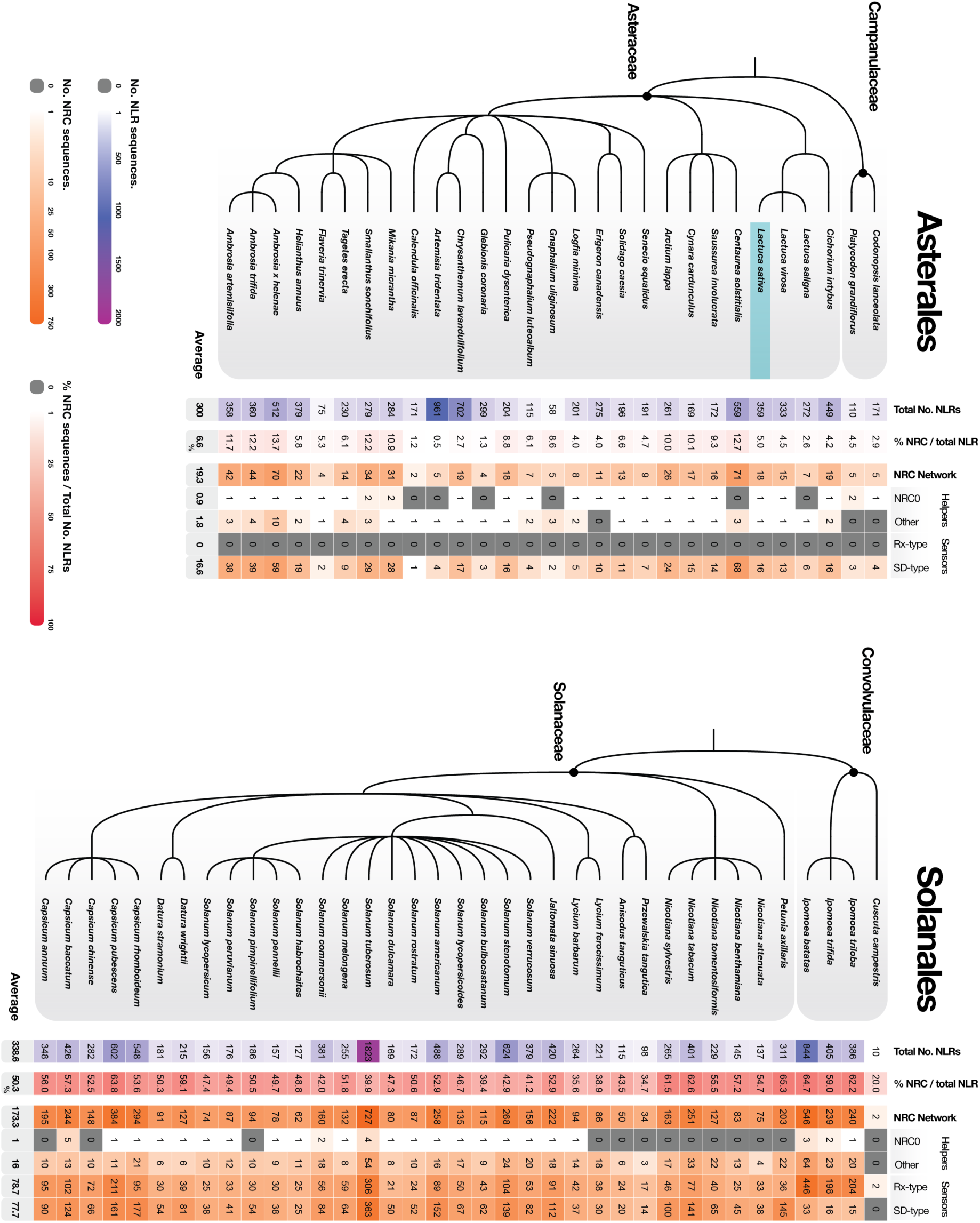
Total number of NLRs, NRC network superclade sequences, NRC0s, other NRC helpers, Rx-type sensors, SD-type sensors, and percentage of NRC superclade sequences out of total number of NLRs in studied Asterales and Solanales species. *Lactuca sativa* (common lettuce) is highlighted in blue.

**Figure S3.**
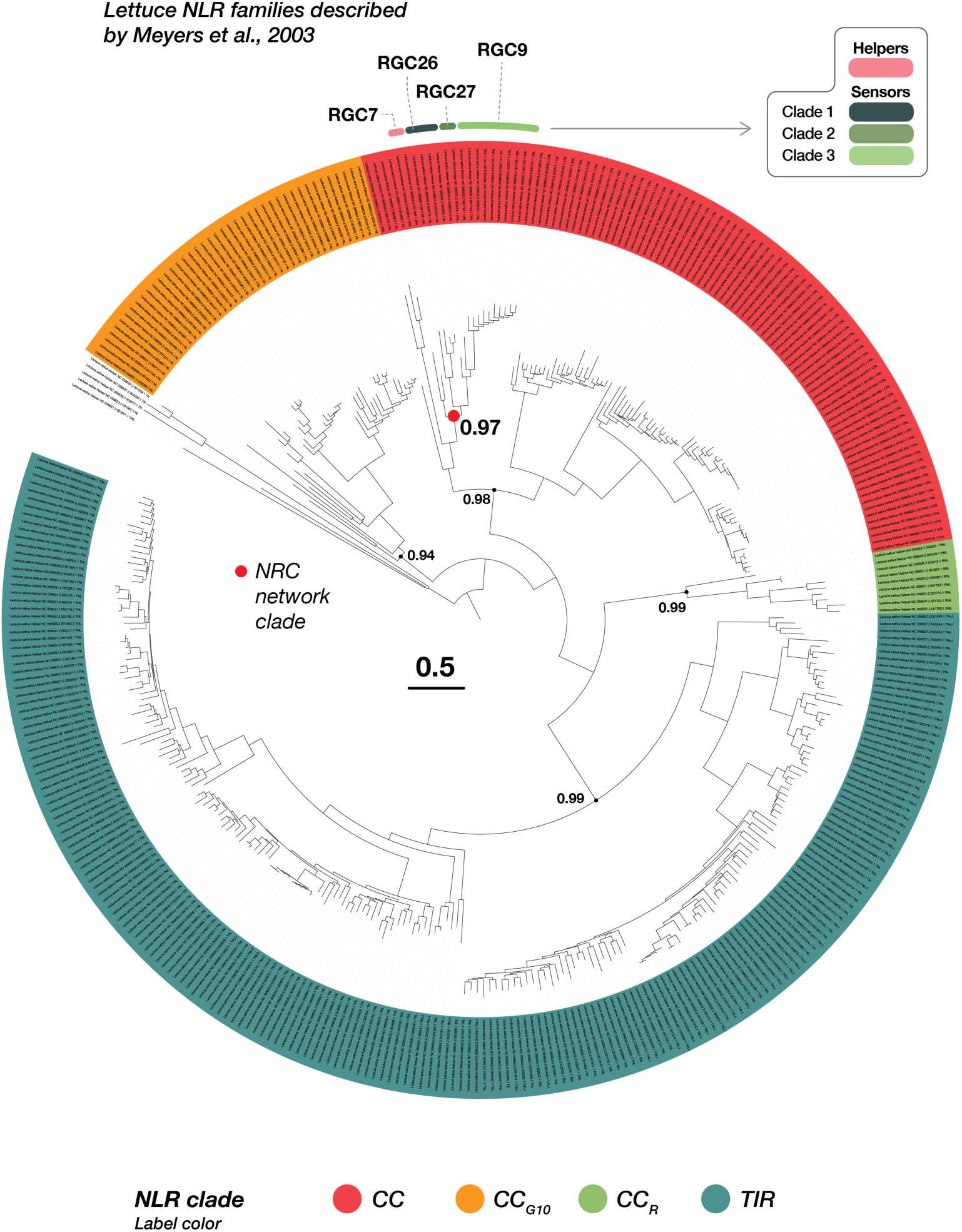
Phylogenetic tree of common lettuce (*Lactuca sativa*) NLRome featuring 359 sequences. Different NLR clades are highlighted in color. Numbers on tree nodes indicate bootstrap values. Phylogenetic tree was made based on NB-ARC domains and rooted at midpoint. NRC network superclade is highlighted with the red dot. NLR family classification by Meyers et al. (1998) is shown for the NRC network superclade (Meyers et al. 1998; Christopoulou et al. 2015).

**Figure S4.**
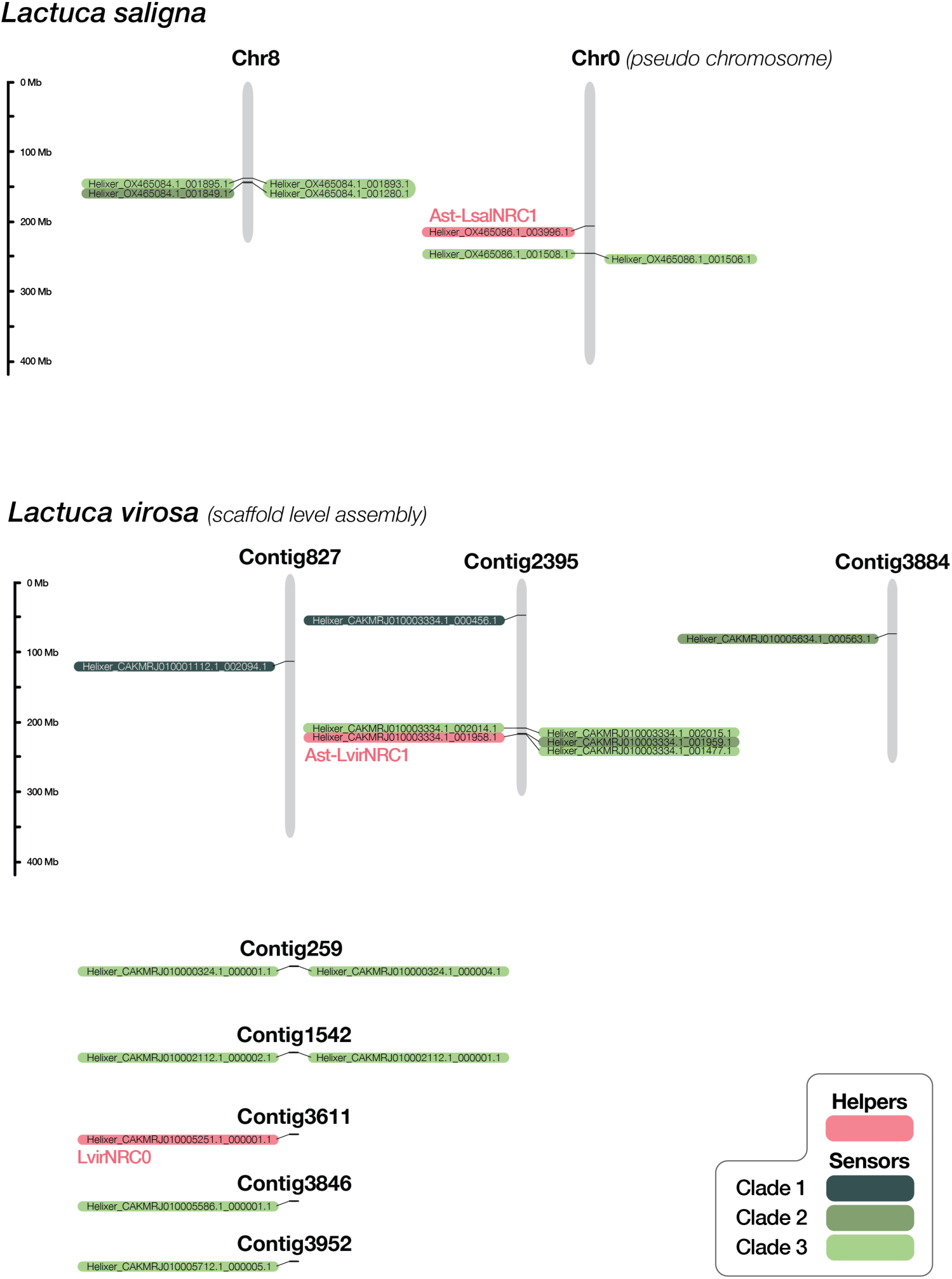
Physical map of *Lactuca saligna* and *Lactuca virosa* NRC superclade sequences.

**Figure S5.**
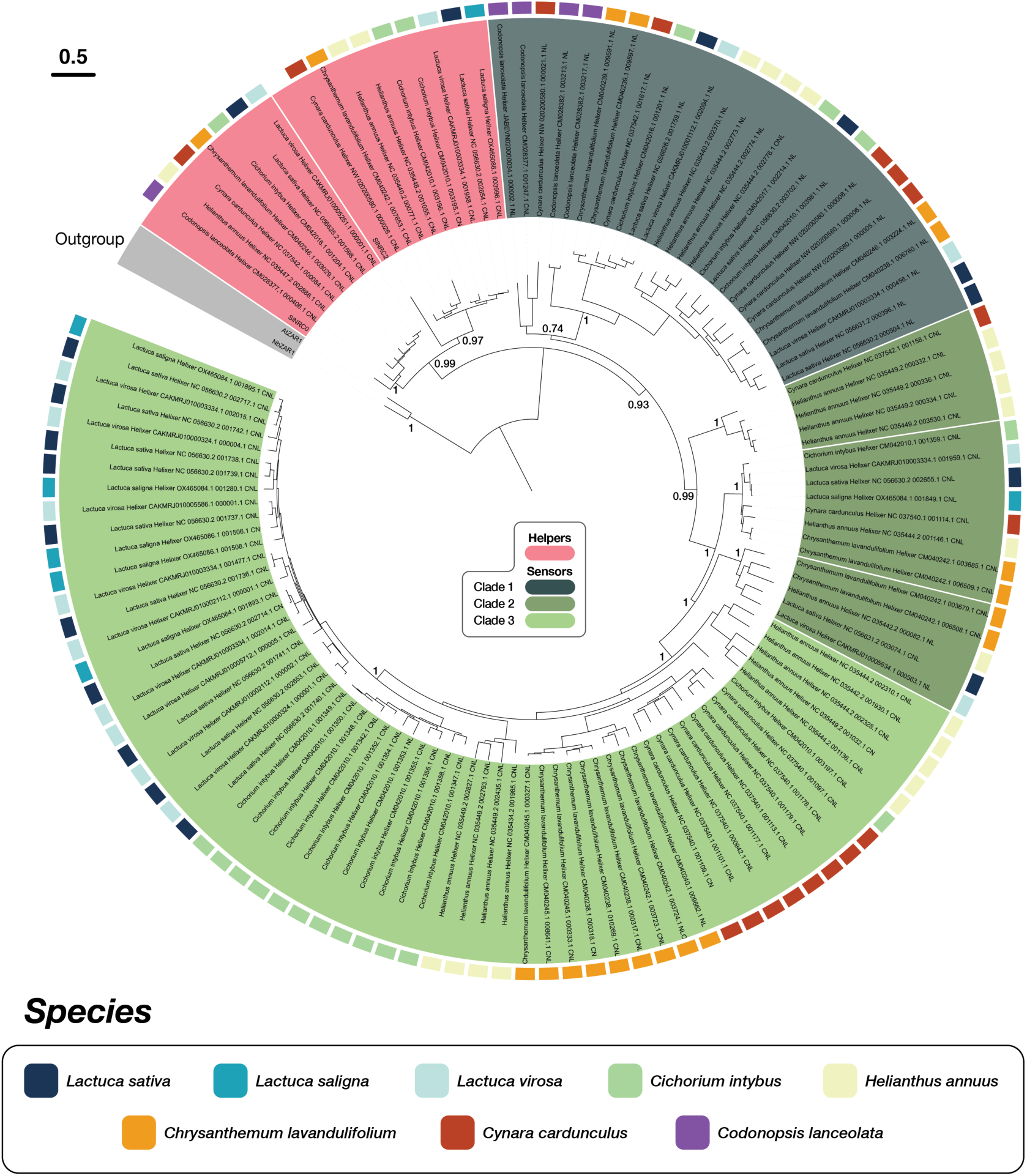
Phylogenetic tree of NRC superclade sequences of Lactuca species (Lactuca sativa, Lactuca saligna, and Lactuca virosa), Codonopsis lanceolata, Helianthus annuus, Cichorium intybus, Chrysanthemum lavandulifolium, and Cynara cardunculus. NRC-S are divided in three phylogenetic categories based on previous classification in the *Lactuca* genus. Numbers on tree nodes indicate bootstrap values. AtZAR1 and NbZAR1 were used as outgroup sequences. SlNRC0 and SlNRC2 were used as reference sequences.

**Figure S6.**
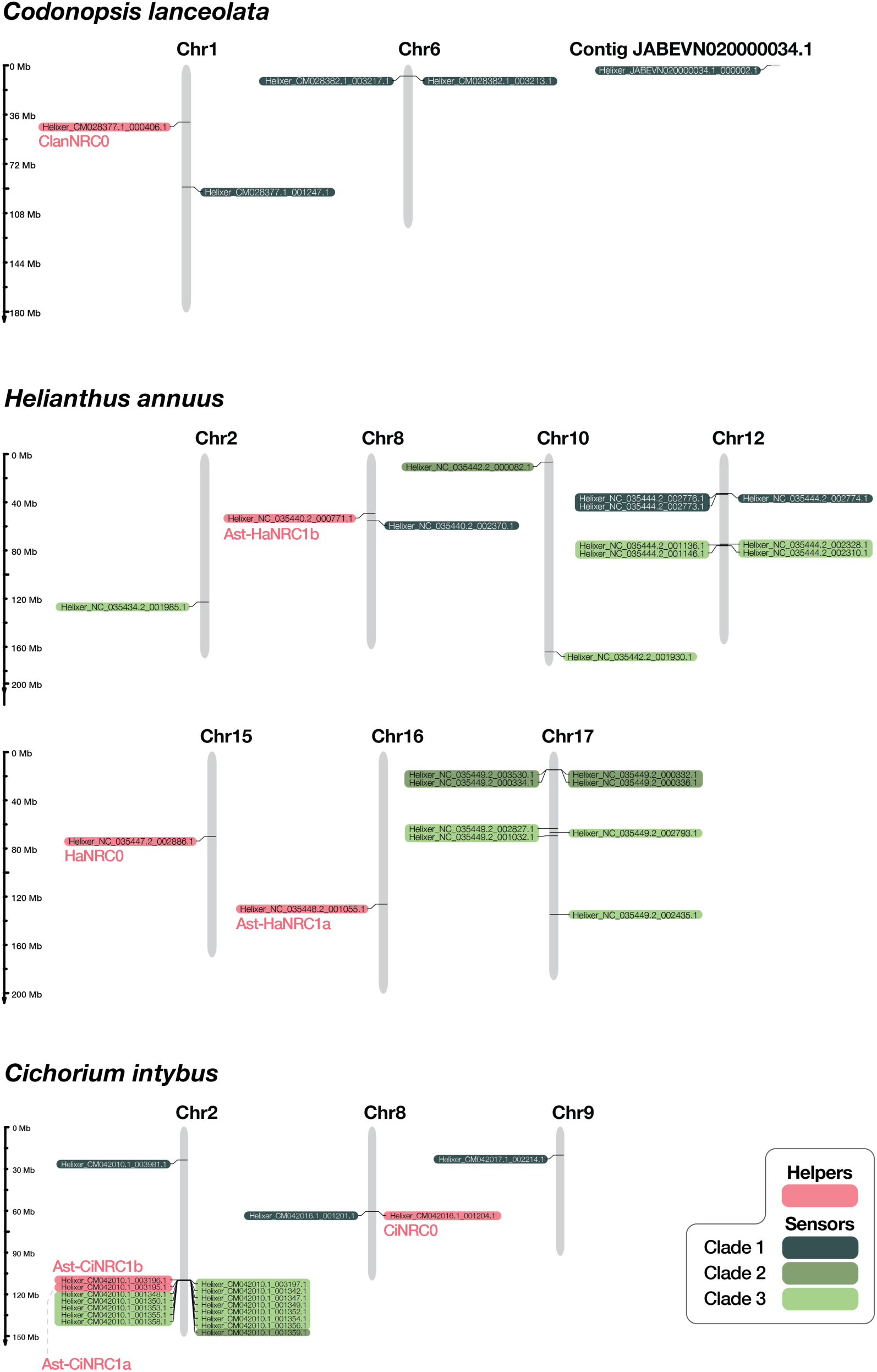
Physical map of *Codonlanceolata*, *Helianthus annuus*, and *Cichorium intybus* NRC superclade sequences.

**Figure S7.**
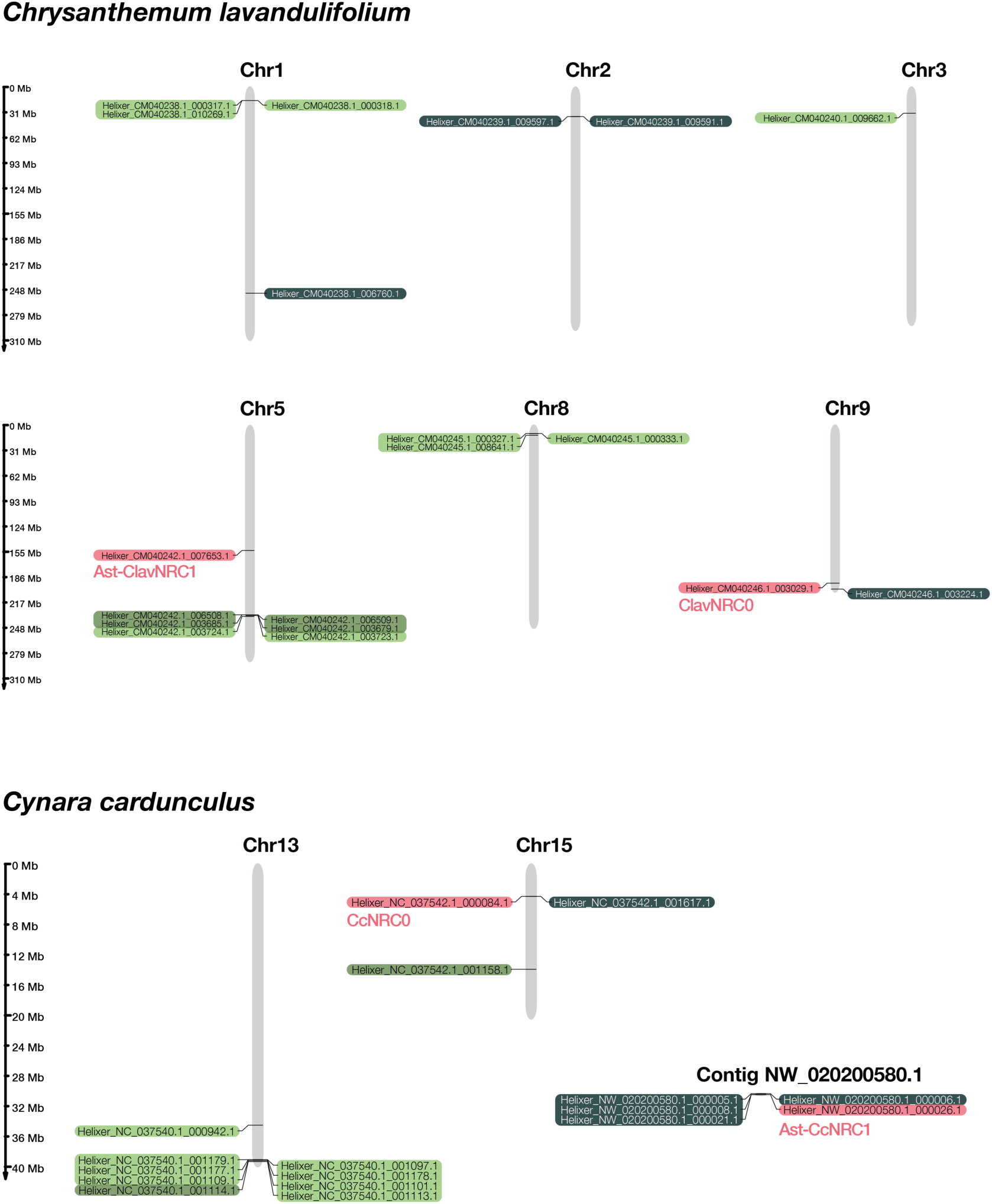
Physical map of *Chrysanthemum lavandulifolium* and *Cynara cardunculus* NRC superclade sequences.

**Figure S8.**
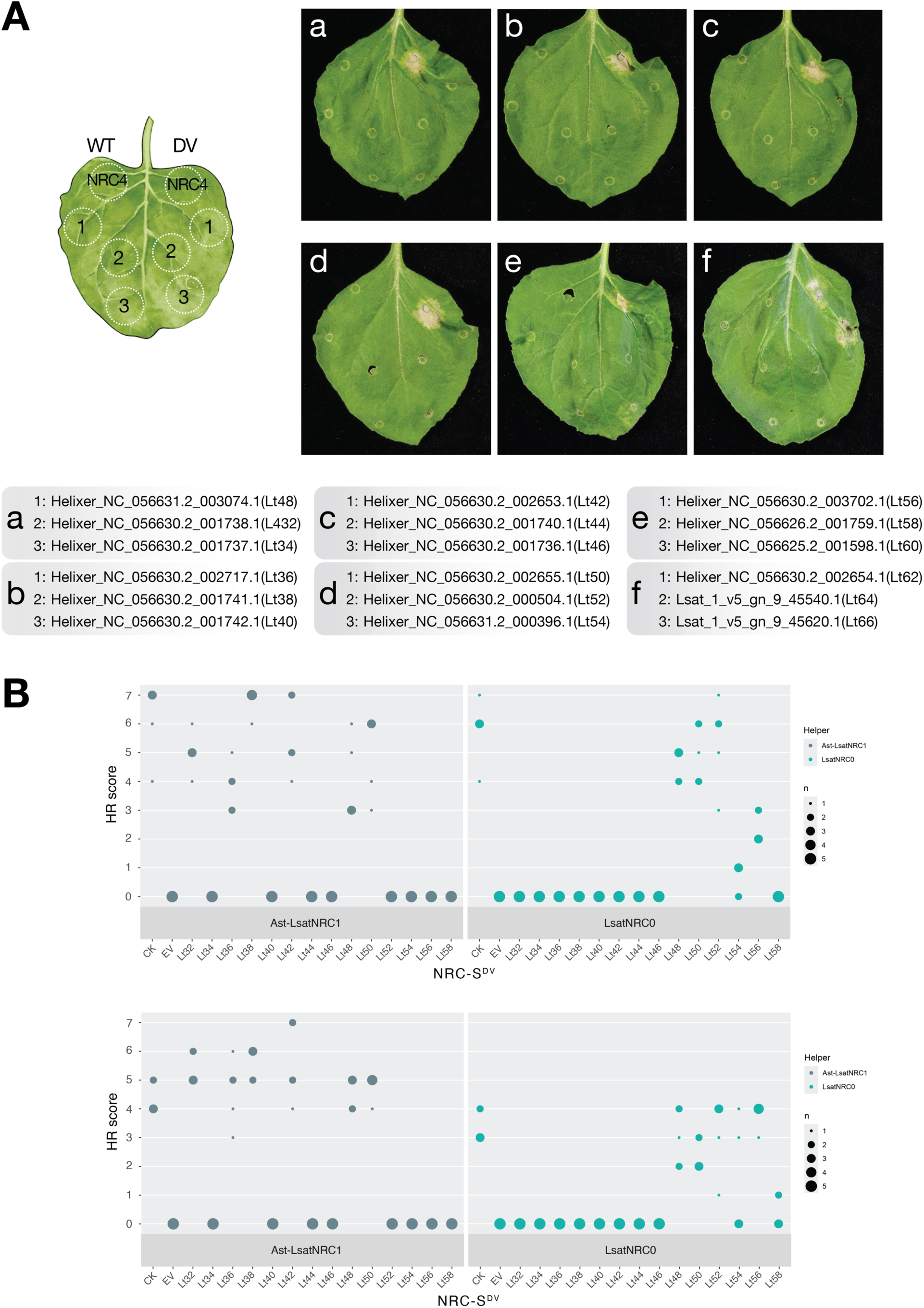
Hypersensitive response (HR) assays of NRC-H and NRC-S. (A) Representative *N. benthamiana* leaves expressing wild-type (WT) or MHD mutated (DV) NRC-H and NRC-S, photographed at 5 days post agroinfiltration. NRC4^WT^ and NRC4^DV^ were infiltrated here as negative and positive controls for HR, respectively. (B) Quantification of co-agroinfiltration HR assays of NRC-H and NRC-S. NRC-S expressed as autoactive mutants with wild-type NRC helpers. CK: NRC4^DV^ infiltrated alone as positive control of HR. EV: NRC-H co-infiltrated with empty vector, as a negative control. Number (n) of data points are reflected as different sized circles. Two biological replicates are displayed as independent panels.

**Figure S9.**
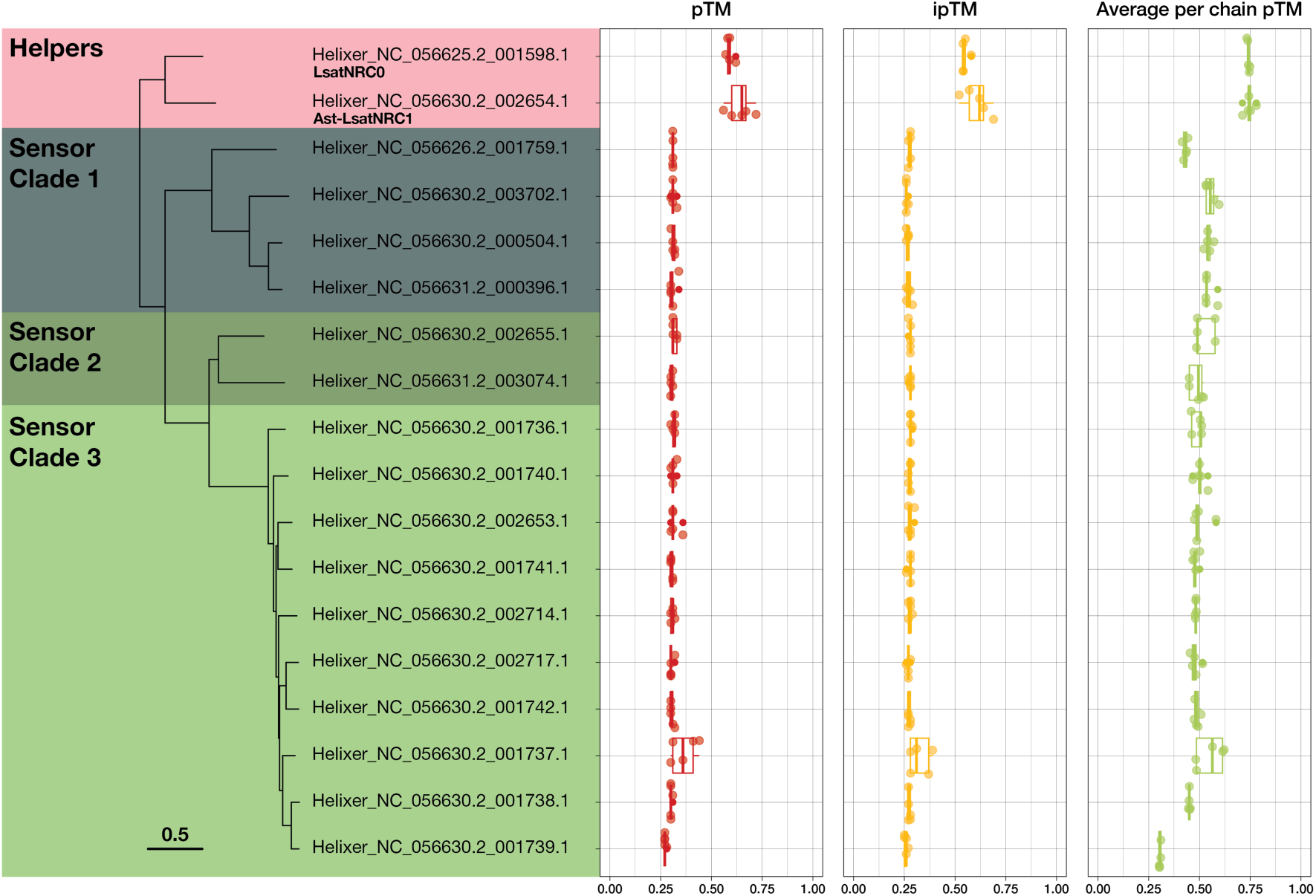
Structural modeling of lettuce NRC-H and NRC-S clade sequences with AlphaFold 3. Simulations included 50 oleic acid molecules to approximate the plasma membrane. Key structural metrics, including pTM (predicted template modeling score for overall structure), ipTM (interface-specific pTM), and per-chain pTM, are plotted for each protein (n=5).

**Figure S10.**
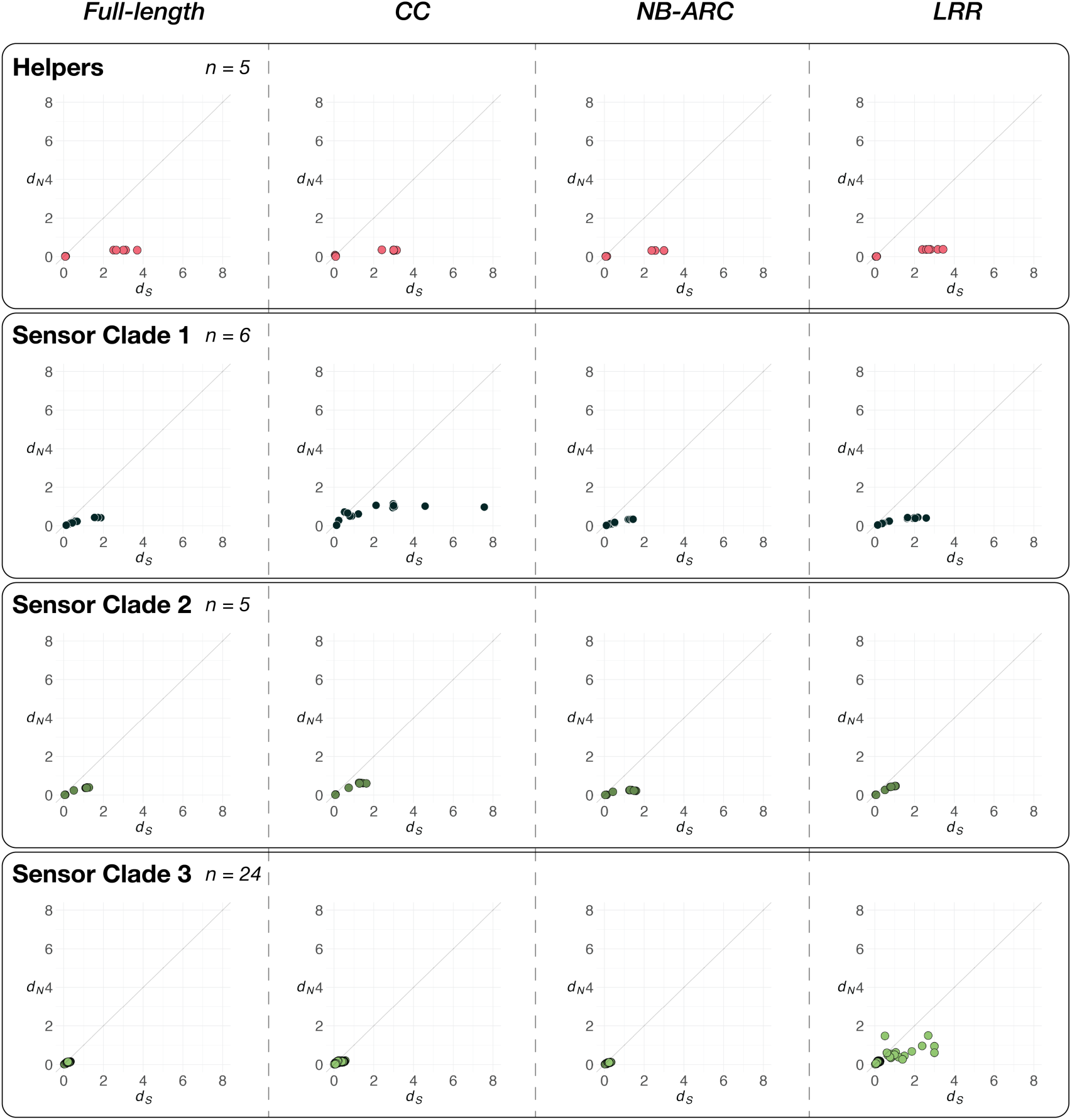
Pairwise *d*_N_ and *d*_S_ values for full-length, and CC, NB-ARC, and LRR domains of NLRs within the *Lactuca* NRC clades. The nonsynonymous (dN) and synonymous (dS) substitution rates were estimated using the approximate method of Nei and Gojobori (1986), implemented in the PAML software (Nei and Gojobori 1986; Yang 1997). A diagonal line represents *d*_N_ = *d*_S_, indicating neutral selection. Points above this line correspond to positive selection (*ω* = *d*_N_ / *d*_S_ > 1).

**Figure S11.**
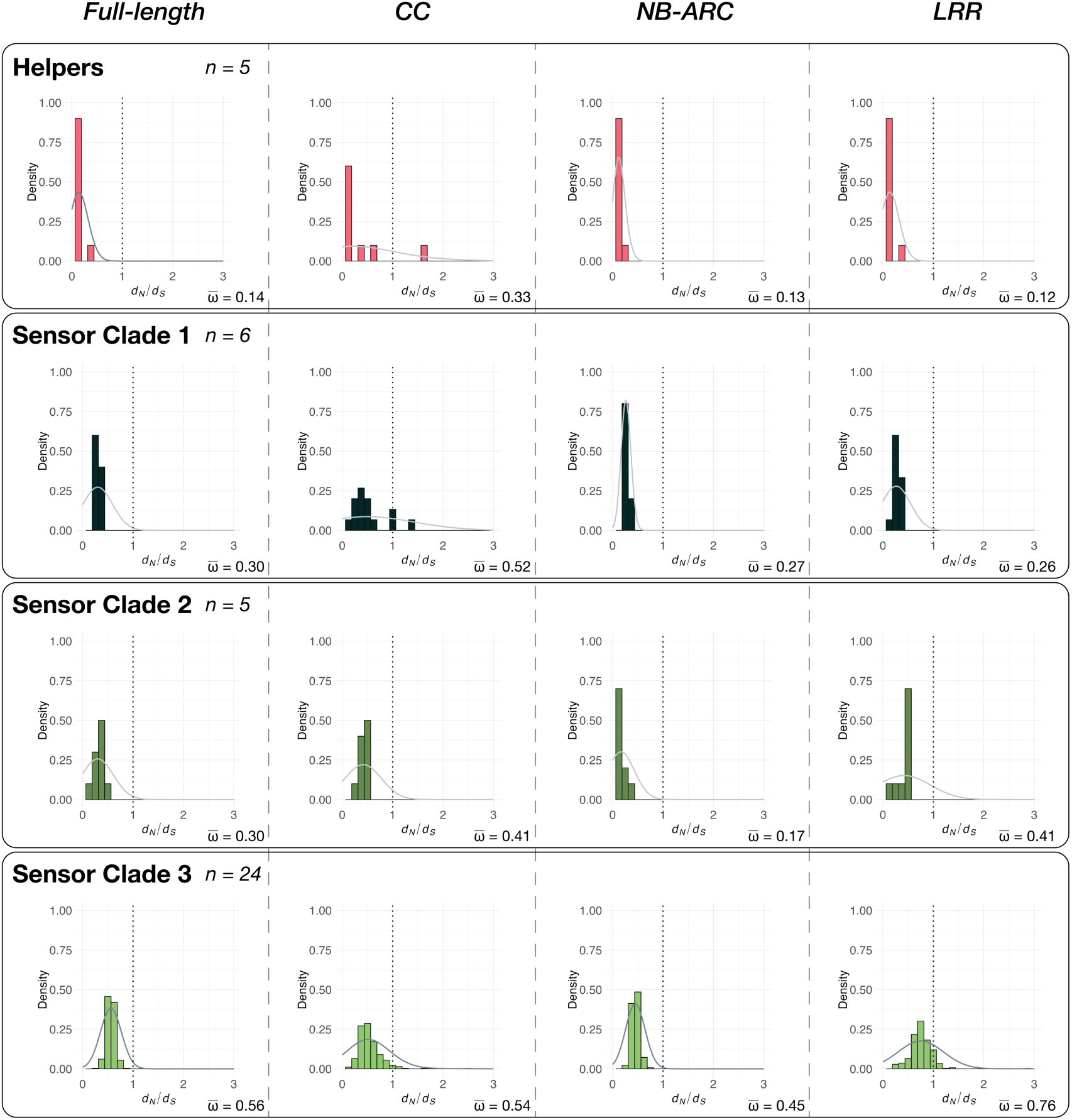
Histogram of pairwise *d*_N_/*d*_S_ ratios for full-length, and CC, NB-ARC, and LRR domains of NLRs within the *Lactuca* NRC clades. The nonsynonymous (dN) to synonymous (dS) substitution rate ratios were estimated using the approximate method of Nei and Gojobori (1986), implemented in the PAML software (Nei and Gojobori 1986; Yang 1997). *d*_N_/*d*_S_ (*ω*) = 1 is highlighted with dotted line. Average *ω* for each graph is displayed at the bottom right corner.

